# Effects of lovastatin on auxin transport and root development in *Arabidopsis thaliana*

**DOI:** 10.64898/2026.02.23.707518

**Authors:** Veronica Giourieva, Christos Tersenidis, Alkiviadis Athanasiadis, Stylianos Poulios, Anna Kouskouveli, Konstantinos Vlachonasios, Emmanuel Panteris, George Komis

**Affiliations:** Department of Botany, School of Biology, Aristotle University of Thessaloniki, 541 24 Thessaloniki, Greece; Natural Products Research Centre of Excellence (NatPro-AUTh), Center of Interdisciplinary Research and Innovation of Aristotle University of Thessaloniki (CIRI-AUTh), 57001 Thessaloniki, Greece

**Keywords:** Lovastatin, auxin gradient, plasma membrane, PINs, cytokinins, FRAP, lateral roots, structural sterols, HMGR

## Abstract

Sterol biosynthesis underlies significant physiological functions in plants, including the production of membrane structural sterols and hormones such as brassinosteroids and cytokinins. Inhibition of sterol biosynthesis has been shown to disrupt multiple aspects of *Arabidopsis thaliana* development. Here, the effects of lovastatin, an inhibitor of HMG-CoA reductase, on root development were investigated, focusing on auxin–cytokinin distribution and transport. Lovastatin inhibited primary root growth, especially cell elongation, in a dose-dependent manner. Additionally, lateral root density was considerably increased and lateral root primordia (LRP) emerged ectopically. In accordance to the above defects, auxin/cytokinin imbalance was recorded by the ectopic presence of the synthetic auxin marker DR5 and a significant decrease of cytokinins, as revealed by depletion of the TCS (two-component signaling) marker. Because auxin distribution appeared disturbed, auxin transport impairment was further examined. Plasma membrane localization of PIN auxin efflux carriers declined significantly, showing additional diffuse cytoplasmic localization in LRP cells. However, the cell-specific localization patterns of several PINs and their abundance at the transcript and protein level appeared unaffected or slightly increased. Fluorescence recovery after photobleaching (FRAP) analysis regarding membrane kinetics of PIN2 revealed altered PIN2 membrane dynamics and transmission electron microscopy (TEM) observations showed structural defects at the plasma membrane-cell wall interface. Together, these results support that sterol biosynthesis is essential for maintaining plasma membrane organization, which, in turn, is key factor for the distribution of hormones that control root development.

**Highlights:** Lovastatin treatment inhibits root growth and causes deregulated formation of lateral roots. Consistently, lovastatin causes altered patterns of auxin distribution relevant to PIN protein mis-localization and decreases cytokinin levels. These changes could be attributed to reduced structural sterols as exemplified from alteration in PIN2 membrane dynamics.

## Introduction

Root development relies on the establishment of highly ordered tissue patterns and a basipetal growth mode starting at the root apical meristem located at the root tip (Dolan *et al*., 1993). Axially, coordinated cell proliferation and differentiation generate distinct developmental zones including meristematic, transition, elongation and differentiation zone (De Nittis *et al*., 2025). Root tissues are arranged in concentric layers, namely the epidermis, cortex, endodermis, pericycle and the centrally located stele containing the vascular tissues (Di Mambro *et al*., 2018).

Proper root growth requires precise communication between cells and tissues over both short and long distances. Auxin plays a central role in this process, governing multiple aspects of root patterning and organization (Benková *et al*., 2003). Directional auxin transport, mediated primarily by PIN-FORMED (PIN) auxin efflux carriers, establishes local maxima underlying primary root growth and lateral root initiation (Lavenus *et al*., 2013; Jourquin *et al*., 2020). Disruption of auxin gradients or transporter localization leads to profound defects in the root system architecture (Benková *et al*., 2003; De Nittis *et al*., 2025).

Lipids, and particularly sterols, have emerged as key regulators of root development through their effects on membrane organization, cell polarity and hormone transport while their deficiency results in post-cytokinesis defects (Souter *et al*., 2002; Willemsen *et al*., 2003; Men *et al*., 2008; Pan *et al*., 2009; Short *et al*., 2018; Boutté and Jaillais, 2020). *Arabidopsis thaliana* mutants defective in sterol biosynthesis before the 24-ethyl/24-methyl sterols checkpoint consistently display altered PIN localization and disturbed auxin distribution in primary roots, underscoring the importance of sterols in maintaining auxin transport routes (Souter *et al*., 2002; Willemsen *et al*., 2003; Men *et al*., 2008; Pan *et al*., 2009).

Sterols comprise about one third of the plasma membrane lipids and are key regulators of membrane fluidity (Grosjean *et al*., 2015; Bahammou *et al*., 2024). In contrast to the animal membrane, plant plasma membrane has a complex sterol composition, including β-sitosterol, stigmasterol and campesterol (Spector and Yorek, 1985; Kierszniowska *et al*., 2009). Sterols are diversely implicated in numerous cellular functions and perturbations of sterol biosynthesis result in developmental defects, including shoot and root growth arrest, vascular mis-patterning, aberrant seed and embryo development, and frequently in lethality (Choe *et al*., 2000; He *et al*., 2003; Schaller, 2003; Kim *et al*., 2005; Carland *et al*., 2010; Cheon *et al*., 2010; Vogel *et al*., 2025).

At cellular level, sterols contribute to the formation of plasma membrane nanodomains that spatially constrain signaling and transport proteins, including auxin carriers, thereby linking membrane structure to developmental signaling outputs. It has been shown that the long-distance auxin transporter ABCB19 depends on sterol-mediated membrane properties for the proper delivery to the plasma membrane, where it functions in concert with PIN1 (Yang *et al*., 2013). Sterol biosynthesis mutants, such as *smt1^orc^* and *cpi1-1,* exhibit defects in endocytosis, cell polarity and hormone signaling (Souter *et al*., 2002; Willemsen *et al*., 2003; Men *et al*., 2008). Although sterols also serve as precursors for brassinosteroids, multiple studies demonstrate that sterols exert brassinosteroid-independent roles in membrane organization and protein trafficking (Willemsen *et al*., 2003; Men *et al*., 2008; Pan *et al*., 2009; Yang *et al*., 2013). However, how total sterol composition affects the localization and dynamics of auxin-related transporters remains elusive. In addition to auxin, cytokinin signaling intersects with membrane trafficking and root zonation, suggesting that sterol-dependent membrane properties may influence multiple hormonal pathways (De Nittis *et al*., 2025; Vogel *et al*., 2025).

Although sterol biosynthesis mutants have been an invaluable tool for elucidating sterol function, their severe developmental phenotypes that often occur complicate the distinction between direct and indirect effects. Pharmacological approaches provide a complementary strategy, allowing temporal and reversible manipulation of sterol biosynthesis. Sterol biosynthesis initiates from acetyl-CoA *via* the cytosolic mevalonate pathway, with 3-hydroxy-3-methylglutaryl-CoA reductase (HMGR) catalyzing a key rate-limiting step (De Vriese *et al*., 2021). HMGR activity can be specifically inhibited by statins, such as lovastatin, which reduces total sterol content and disrupts multiple developmental processes in *A. thaliana* (Suzuki *et al*., 2004; Kobayashi *et al*., 2007; Vogel *et al*., 2025).

As previously shown, lovastatin inhibits primary root elongation, arrests true leaf development and induces dwarfism (Baskin and Bivens, 1995; Nagata *et al*., 2002; Kobayashi *et al*., 2007). These effects are distinguished from those caused by brassinosteroid deficiencies, as exogenous supplementation with brassinolide does not rescue the sterol deficient growth defects (Suzuki *et al*., 2004).

Despite genetic analyses of sterol biosynthesis, the effects of pharmacological inhibition of total sterol production on root system architecture remain elusive. Therefore, the effect of lovastatin on the development of root architecture was investigated. Lovastatin reversibly affected primary root development in a dose-dependent manner. Inhibition of sterol biosynthesis distrurbed the auxin-cytokinin equilibrium, resulting in the shrinkage of primary root developmental zones and mis-regulation of lateral root emergence, mis-localization of major PINs, probably as a result of defective membrane trafficking. Together, these findings highlight the critical role of sterols in maintaining plasma membrane organization and auxin transporter localization during early root system development. Given the conserved function of sterols in eukaryotic membrane organization, these mechanisms are likely to extend beyond *A. thaliana* to other plant species.

## Materials and methods

### Plant material and growth conditions

*Arabidopsis thaliana* (L) Heynh, ecotype Columbia-0 (Col-0) and reporter lines in the same ecotype background, listed in Supplementary Table S1, were grown in a growth chamber at 21-22°C under long day conditions (16h light/8 h dark) and light intensity 120 µmol m^− 2^ s^− 1^. Prior to plating, seeds were surface sterilized with 30% (v/v) sodium hypochlorite supplemented with 0.02% (v/v) Triton-X100 and stratified at 4°C for 2 d in the dark. Half strength Murashige and Skoog (1/2 MS; Duchefa, Haarlem, the Netherlands) salt medium supplemented with 2% (w/v) sucrose (Applichem, Darmstadt, Germany) and 1% (w/v) phytoagar (Duchefa, Haarlem, the Netherlands), pH 5.4-5.7, was used for seed plating. Seedlings were grown vertically in Petri dishes.

### Chemical treatment

For the treatments with the sterol biosynthesis inhibitor lovastatin (Cat no PHR1285, Merck, Darmstadt, Germany) a 1 mM stock solution was prepared in anhydrous DMSO and stored in -20°C. For the dose response treatments, additional stocks of 0.01 mM, 0.05 mM, 0.075 mM, 0.1 mM, 0.3 mM and 1 mM were prepared and added directly to autoclaved media, appropriately diluted in 1/2 MS at a range of final working concentration of 0.01 μΜ, 0.05 μΜ, 0.075 μΜ, 0.1 μΜ, 0.3 μΜ and 1 μΜ, respectively. Seeds were grown directly in Petri dishes containing the inhibitor. For the recovery experiments, seeds were initially grown on medium containing the appropriate concentration of lovastatin and, after 4 days of treatment, they were transferred to 1/2 MS medium devoid of lovastatin. The potential of recovery was scored after 6 days. Mock samples were grown in 1/2 M/S medium supplemented with 0.1% (v/v) DMSO.

### Root growth and morphology characterization and quantitative analysis

For the characterization of hypocotyl and root growth, seeds were grown for 5 d after germination (DAG) in 0.01 μΜ, 0.05 μΜ, 0.075 μΜ, 0.1 μΜ, 0.3 μΜ and 1 μΜ lovastatin in solid 1/2 MS and subsequently documented. Images of hypocotyls and roots were acquired with a Stereo Discovery V8 stereomicroscope (Carl Zeiss, Berlin, Germany) equipped with Axiocam ERc 5s camera. Root growth rates were acquired after the second day of germination over a period of 8 d. Hypocotyl length and width, root length and distance of the first bulging trichoblast from root apex were measured at 10 DAG. The number of lateral root primordia (LRP), lateral roots and length of the first epidermal cell with visible root hair bulge (LEH) (Le *et al*., 2001) were calculated from differential interference (DIC) images obtained under AxioImager.Z2 light microscope equipped with AxioCam MRc 5 (Carl Zeiss, Berlin, Germany), by Axiovision rel 4.8 software from 10 d-old seedlings. The stages of LRPs were classified according to Malamy and Benfey (1997). For the recovery assay, seeds were germinated in 0.01 μΜ, 0.1 μΜ, 0.3 μΜ and 1 μΜ lovastatin, transferred to lovastatin-free 1/2 MS medium and root growth was measured for 5 d after the transfer. Statistical analysis was performed with Graphpad Prism 8.0.1 software using one or two-way analysis of variance (ANOVA) with multiple comparison (Tukey’s or Dunn’s test depending on the samples distribution). Kolmogorov-Smirnov test was used for evaluation of Gaussian distribution. Imaging and processing settings were the same for both treated and untreated seedlings.

### Root clearing and cell wall fluorostaining

For tissue patterning analysis, roots of the reporter lines were counterstained by propidium iodide (20 μg/ml in PBS) for short exposure duration to lovastatin, while for longer exposure (seedlings older than 7 d and LRPs) they were fluorostained with Calcofluor White M2R (Cat no 18909, Sigma-Aldrich, Darmstadt, Germany). Calcofluor White staining was performed according to Ursache *et al*. (2018) with minor modifications. Specifically, samples were stained overnight with 0.05% (w/v) Calcofluor White in clearing solution (“ClearSee”: 10% w/v xylitol, 15% w/v sodium deoxycholate and 25% w/v urea). Images were acquired by confocal laser scanning microscopy (CLSM) with a ZeissObsderver.Z1 inverted microscope equipped with LSM 780 module (Carl Zeiss, Germany). A Plan-Apochromat 40x/1.3 NA oil immersion objective was used and analysis was performed with Zeiss Zen 3.13 (Zen lite, Blue version). Propidium iodide and Calcofluor White were excited at 543 nm (65% laser intensity) and 405 nm (8-12% laser intensity) and detected at 595-632 nm and 410-524 nm, respectively. Calcofluor White staining was also used to determine the length of root meristem (number of cortex cells between the quiescent center and the first elongating cortex cell). Length of root meristem was determined according to Perilli and Sabatini (2010).

### 3D root image acquisition

For 3D imaging of the Calcofluor White stained roots, z-stacks were acquired with LSM780 confocal microscope, using the same illumination settings as described above, with pinhole at 7.5 AU/90 μm and z-steps of 1.16 μm. Image pre-processing was done in ZEN 3.11 software, using the orthogonal projection capabilities. Orthogonal view from different image planes (x/y, x/z or y/z) were used to detect the columella and stele initials, to analyse the quiescent centre (QC)organisation and the radial root patterning. Cell numbers were counted using the Fiji plugin “Cell counter” (Schindelin *et al*., 2012).

### Light and transmission electron microscopy (TEM)

Seedling roots of mock seedlings and those treated with 1μM lovastatin for 4, 5 or 7 d were prepared for light microscopy and TEM according to Panteris *et al*. (2021). In brief, 2-3 mm apical root segments were fixed in 3% v/v glutaraldehyde in 50 mM sodium cacodylate buffer (pH 7) for 4 h and post-fixed in 1% w/v osmium tetroxide in the same buffer for 3 h. Afterwards, the samples were dehydrated in a graded acetone series and subsequently with anhydrous propylene oxide 2 x 30 min and finally embedded in Spurr’s resin. Semi-thin sections (1.5-2 μm) were obtained with a Reichert-Jung Ultracut E ultramicrotome (Reichert-Jung Optical Company, Vienna, Austria), stained with 1% (w/v) toluidine blue O in 1% (w/v) aqueous borax and examined under an AxioImager.Z2 light microscope. Ultrathin sections (70-90 nm) were cut with the above microtome with a diamond knife, collected on copper grids and double stained with uranyl acetate and Reynold’s lead citrate. The specimens were observed with a JEOL JEM 1011 TEM (TEM, JEOL, Ltd., Tokyo, Japan) at 80 kV. Electron micrographs were acquired with a Gatan ES500 W camera (Gatan Inc., Tokyo, Japan) and the Digital Micrograph 3.11.2 software (Gatan Inc.).

### RNA extraction and qPCR

For the RT-qPCR expression analysis, roots from 4-d-old Col-0 plants, either mock or lovastatin-treated, were collected and flash-frozen in liquid nitrogen. The frozen tissue was preserved at −80 °C. RNA extraction was performed using the Nucleospin® RNA Plant kit (Macherey-Nagel, Duren, Germany). RNA quality and quantity were assessed using 1.5% agarose gel electrophoresis and spectrophotometrically using NanoDrop 2000 (Thermo Fischer Scientific, Waltham, MA, USA). The PrimeScript^TM^ 1st strand cDNA synthesis kit (Takara, Shiga, Japan) was used for reverse transcription. In three independent biological repeats, reverse transcription was performed using 130ng of total RNA. Quantitative reverse-transcription polymerase chain reactions (RT-qPCRs) were prepared with the Luna® Universal qPCR Master Mix (New England Biolabs, Ipswich, MA, USA) using the ABI StepOnePlus™ system (Applied Biosystems, Foster City, CA, USA). Three technical repeats were run for each sample. The At4g26410 gene (RHIP1) were used as endogenous control (Suppl Table S2). Data were analysed with the ΔΔCt method using StepOne Software 2.1 (Applied Biosystems, Foster City, CA, USA). Statistical analysis was performed using the GraphPad Prism8.0.1 software.

### Protein extraction, SDS-PAGE and immunoblot analysis

Four-day-old seedlings of reporter lines *PIN1::PIN1-GFP*, *PIN2::PIN2-GFP*, *PIN3::PIN3-GFP* and *PIN4::PIN4-GFP* grown on 1 μM lovastatin or 0.1% (v/v) DMSO were used. Total proteins were extracted from roots using RIPA buffer (50 mM Tris-HCl pH 7.4, 150 mM KCl, 5 mM EGTA, 0.1% SDS, 0.5% sodium deoxycholate, 100 mM dithiothreitol) supplemented with GRS protease inhibitor cocktail (Grisp Research Solutions, Portugal). After quantification of protein content with Bradford assay or measuring absorbance at 280 nm, protein extracts were diluted with Laemmli loading buffer (50 mM Tris-HCl pH 6.8, 2% (w/v) SDS, 0.1% (w/v) bromophenol blue and 2.5% (v/v) *β*-mercaptoethanol, (Laemmli, 1970) and denatured for 3 min at 80°C. Samples were separated in 10% (w/v) SDS/polyacrylamide gels and transferred to 0.45 μm pore size polyvinylidene fluoride membranes (Ultracruz sc-3723, Santa Cruz Biotechnology, Texas, USA) by electroblotting. Total proteins were visualized with silver staining according to Blum et al. (1987). The membranes were incubated with 1:1000 monoclonal rabbit antibody raised against GFP (#2956, Cell signalling technology, Danvers, MA, USA) and 1:2000 monoclonal mouse anti-actin antibody (A0480, Sigma Aldrich, St Louis, MO, USA). As secondary antibodies, horseradish perodixase-conjugated goat anti-rabbit (1:2000) and rabbit anti-mouse (1:2000) were used (Cell signalling technology, Danvers, MA, USA). Blots were developed using the Agrisera ECL Light (Agrisera AB, Sweden) according to manufacturer’s instructions and visualized with the Invitrogen iBright FL1500 documentation system (Invitrogen, Waltham, MA, USA). Analysis of density of the bands was carried out with ImageJ (Schneider *et al*., 2012). The Coomassie-stained and actin immunodetected samples were used as internal loading controls for the protein expression. Three biological replicates and at least two technical replicates were used for the PINs quantification.

### Membrane dynamics-Fluorescence recovery after photobleaching (FRAP)

Four-day-old *PIN2::PIN2-GFP* seedlings were used for the FRAP analysis. Seedlings were mounted in liquid 1/2 M/S (pH 5.4-5.7), 2% (w/v) sucrose and 1 μM lovastatin or 0.1% (v/v) DMSO on a 24 x 50 mm coverslip and fixed with an additional 24 x 24 mm coverslip to avoid excessive pressure to the samples. Cotyledons remained outside of the coverslips to prevent uneven mounting and sample drift during analysis. FRAP was performed with LSM 780 and 488 nm argon laser, using the oil immersion 40x/1.3 NA objective and pinhole size 160. GFP signal was detected between 493 nm and 598 nm. For the FRAP experiments the Zen 2011 software with active bleaching tool group was employed and three pre-bleach images were acquired. Samples were photobleached with 100% of 488 nm laser until fluorescent intensity dropped to 50% and 25 post-bleach images were acquired with a 6 s interval. Pre-bleach and post-bleach scans were performed with 4% laser power. For all fluorescence intensity analysis (normalization, graph generation, calculation of mobile/immobile fraction, t1/2 and standard error calculation) of FRAP experiments, intensities of bleached and unbleached regions were extracted as *.txt files and subjected to the in-house standalone software FRAPedia 1.0.0 designed to analyze FRAP datasets using Python 3.11. At least 6 seedlings with at least four regions were used from each condition. Analysis of normal distribution was performed with Kolmogorov-Smirnov test combined with t-test.

### Statistical analysis

For the quantitative analysis of root development aspects, one or two-way analysis of variance (ANOVA) with multiple comparison post-hoc (Tukey s or Dunn’s test depending on the samples distribution) was applied. Kolmogorov-Smirnov test was used for normality distribution. At least three biological and three technical replicates were used unless otherwise stated. For FRAP experiments at least four seedlings and four regions per seedling were evaluated. Statistical analysis conducted with Microsoft Office365 Excel or GraphPad Prism 8.0.1.

## Results

### Lovastatin treatment reversibly inhibited root growth in a dose-dependent manner

Primary root growth was followed over a broad range of lovastatin concentrations (0.01μM - 1μM; Fig.1, Supplementary Fig. S1, Supplementary Fig. S2) with the effective range being restricted to 0.1 μM – 1μM.

**Figure 1:**
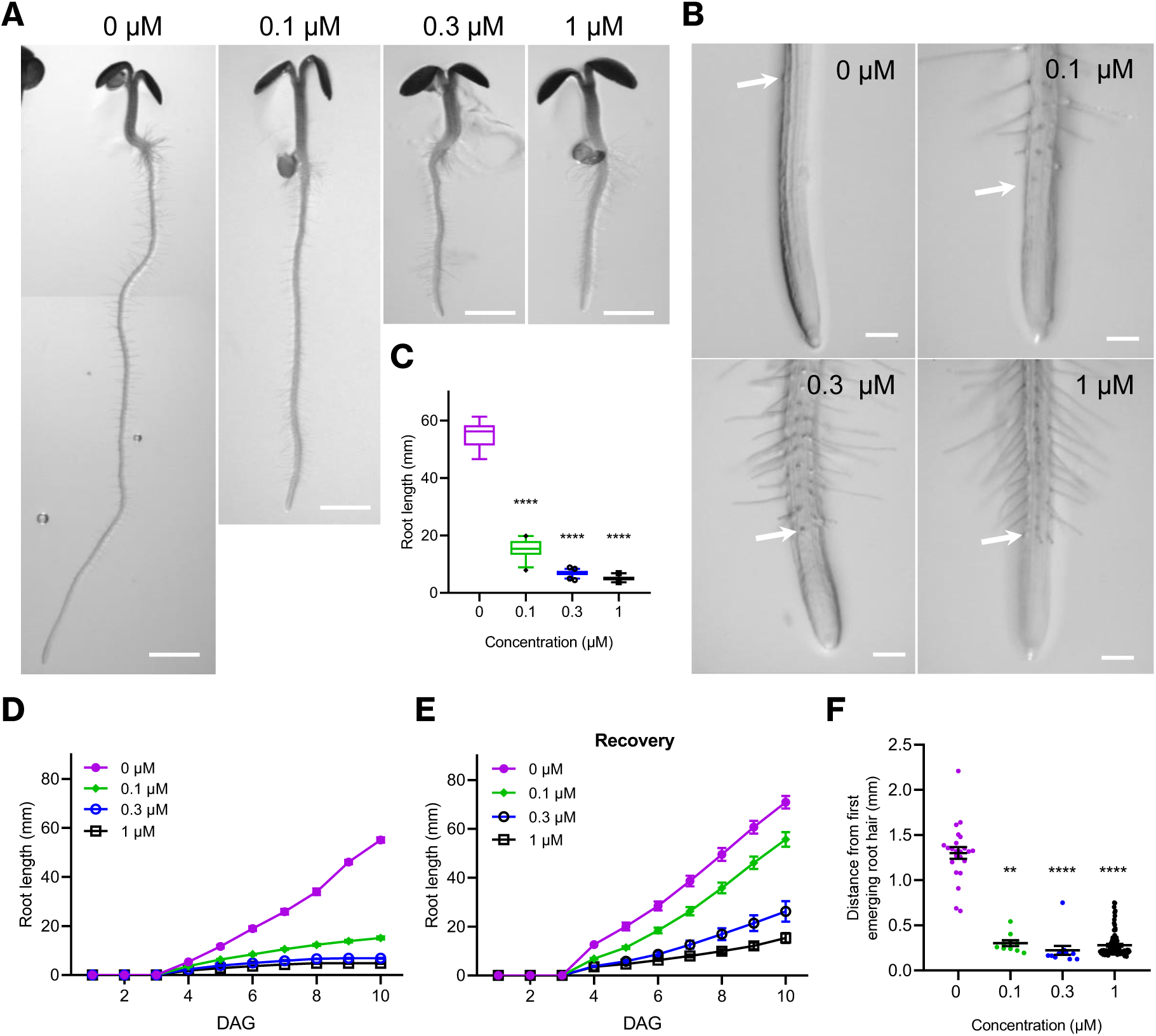
Lovastatin reversibly inhibits root growth after 4 DAG. (A) Five-day-old seedlings grown in medium with 0, 0.1, 0.3 and 1 μM lovastatin. (B) Respective root apex of lovastatin-treated seedlings. (C) Root length of 7-d-old seedlings grown in the presence of different lovastatin concentration. Kruskal’s Wallis test combined with Dunn’s multiple comparison. For 0 μM N=17, 0.1 μΜ: N=26, 0.3 μM: N=42, 1 μM: N=21. Comparison was made against 0 μΜ (mock). Bars represent 5-95% of sample distribution and outliers are shown. (D) Growth rate of lovastatin-treated seedlings for 10 d. 0 μM: N=18, 0.1 μM: N=27, 0.3 μM: N=22, 1 μM: N=43. (E): Recovery of 4 d lovastatin-treated seedlings with 0 μM (Ν=16), 0.1 μM (Ν=22), 0.3 μM (Ν=24) and 1 μΜ (Ν=24). (F): Distance of the first emerging root hair decreased significantly (compared to 0 μΜ (Ν=23), 0.1 μM (N=10), 0.3 μM (N=12) and 1 μM (N=86). ** *p*<0.001, **** *p*<0.0001. Scale bar: (A) 1 mm, (B): 0.1 mm. N: number of different plants. At 3 biological replicates were used.

*A. thaliana* seeds germinate normally and seedlings exhibit uninterrupted growth for *ca.* 3 DAG on lovastatin-supplemented medium. Afterwards, root (Fig. 1A) and hypocotyl (Supplementary Fig. S2) elongation slows down with lovastatin treatment, in a dose dependent manner, resulting in stunted growth and accompanying developmental defects (Fig. 1, Supplementary Fig. S1, Supplementary Fig. S2).

Subsequently, lovastatin treatment caused significant decrease in root elongation rate over a period of 10 days (Fig.1A, B, C, D, Table 1, Table 2). Primary roots treated with lovastatin for 4-5 d (for 0.3 μM and 1 μΜ) were considerably shorter, compared to those of mock seedlings (Fig1 A, B, Table 1), while root elongation rate was compromised at all concentrations above 0.1 μM (Fig 1C, D, Table 2). At concentrations ≥ 0.3 μM root growth was fully arrested (Fig 1 A-D, Fig. 1F, Supplementary Fig. S1, Table 1, Table 2). The effects of lovastatin on root growth were partially reversible, as proven for lovastatin-treated seedlings subsequently transferred to control medium at the end of the treatment (Fig. 1 E, Supplementary Fig. S3). Recovery was almost complete after treatment with the lowest effective lovastatin treatment (0.1 μM) and only partial after treatment with 0.3 μM and 1 μM (Supplementary Fig. S3, Table 1, Table 2).

**Table 1:**
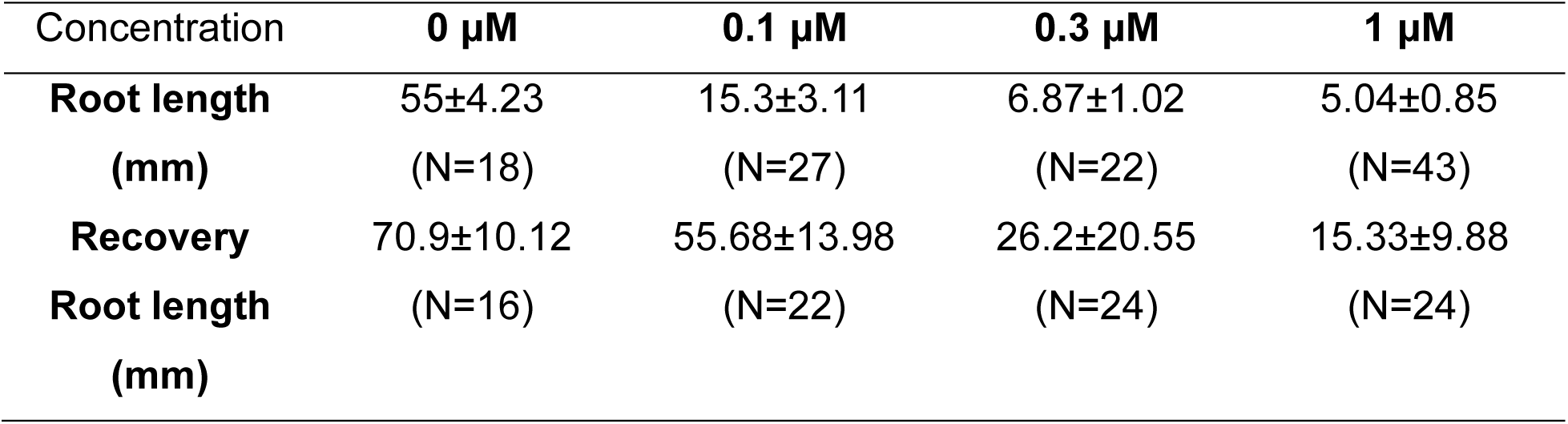
Root length of mock and lovastatin-treated seedlings 10 DAG. Recovery root length refers to the seedlings grown in mock or lovastatin-supplemented media and transferred to control media 4 DAG.

**Table 2:**
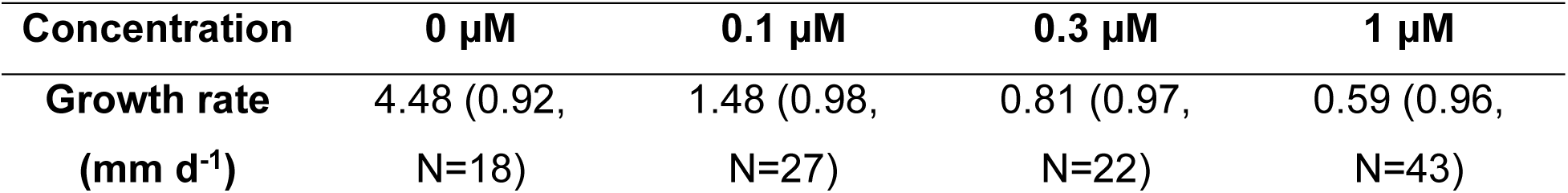
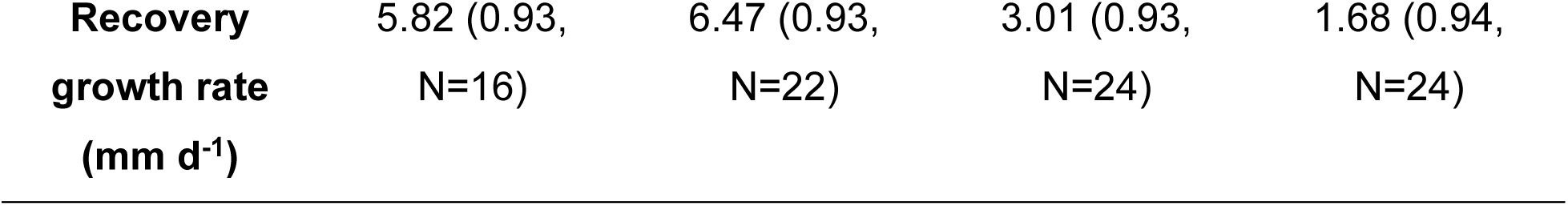
Root growth rate of seedlings, mock or treated with 0.1 μM, 0.3 μM and 1 μM lovastatin for 10 d. Recovery growth rate corresponds to recovery assays. Numbers in brackets R^2^, N: number of seedlings used. At least 3 biological replicates were used.

In accordance with lovastatin-induced root growth inhibition, the distance between the first emerging root hair (first trichoblast with discernible bulge) and root tip gradually decreased, while root hair density increased, dose-dependently (Fig 1B, F, Table 3). This could be attributed to either a premature differentiation of epidermal cells and/or shrinkage of root developmental zones. Apart from the root, hypocotyl growth was also significantly reduced compared to the untreated seedlings (Supplementary Fig. S2). Emergence of leaves from the shoot apical meristem was notably inhibited by lovastatin (Supplementary Fig. S2). As in roots, the effects of lovastatin in hypocotyl development were reversible upon transfer of seedlings to lovastatin-free medium, allowing, among others, the emergence of leaves from arrested primordia in the shoot apical meristem (Supplementary Fig. S3).

**Table 3.** Distance of first emerging root hair from root tip decreased significantly. Distance of root tip from the first emerging root hair of seedlings 7 DAG. Kruskal-Wallis test and Dunn’s multiple comparison tests were used. **p<0.01, ****p<0.0001.

| **Concentration (μM)** | **Distance of root tip from the first emerging**<br>**root hair (mm)** |
| --- | --- |
| **0** | 1.3±0.065 (N=23) |
| **0.1** | 0.3±0.03 (N=10, **) |
| **0.3** | 0.22±0.05 (N=12, ****) |
| **1** | 0.28±0.015 (N=86, ****) |

### Treatment with lovastatin affected the length of root developmental zones but not radial root organization

Following the general effects of lovastatin on primary root growth, a more detailed detection of the origins of root defects was sought. Therefore, the effects of lovastatin on cell shape, size and tissue patterning were examined in appropriately cleared roots stained with Calcofluor White to delineate cell borders (Ursache *et al*., 2018). Root developmental zones were classified according to Perilli and Sabatini (2010) and their length and cell patterning were determined according to the number of cortex cells per developmental zone (Fig. 2, Supplementary Fig. S4). Specifically, for the determination of meristem size, cortex cells from QC to the first elongating cell were analyzed (Fig. 2A).

**Figure 2:**
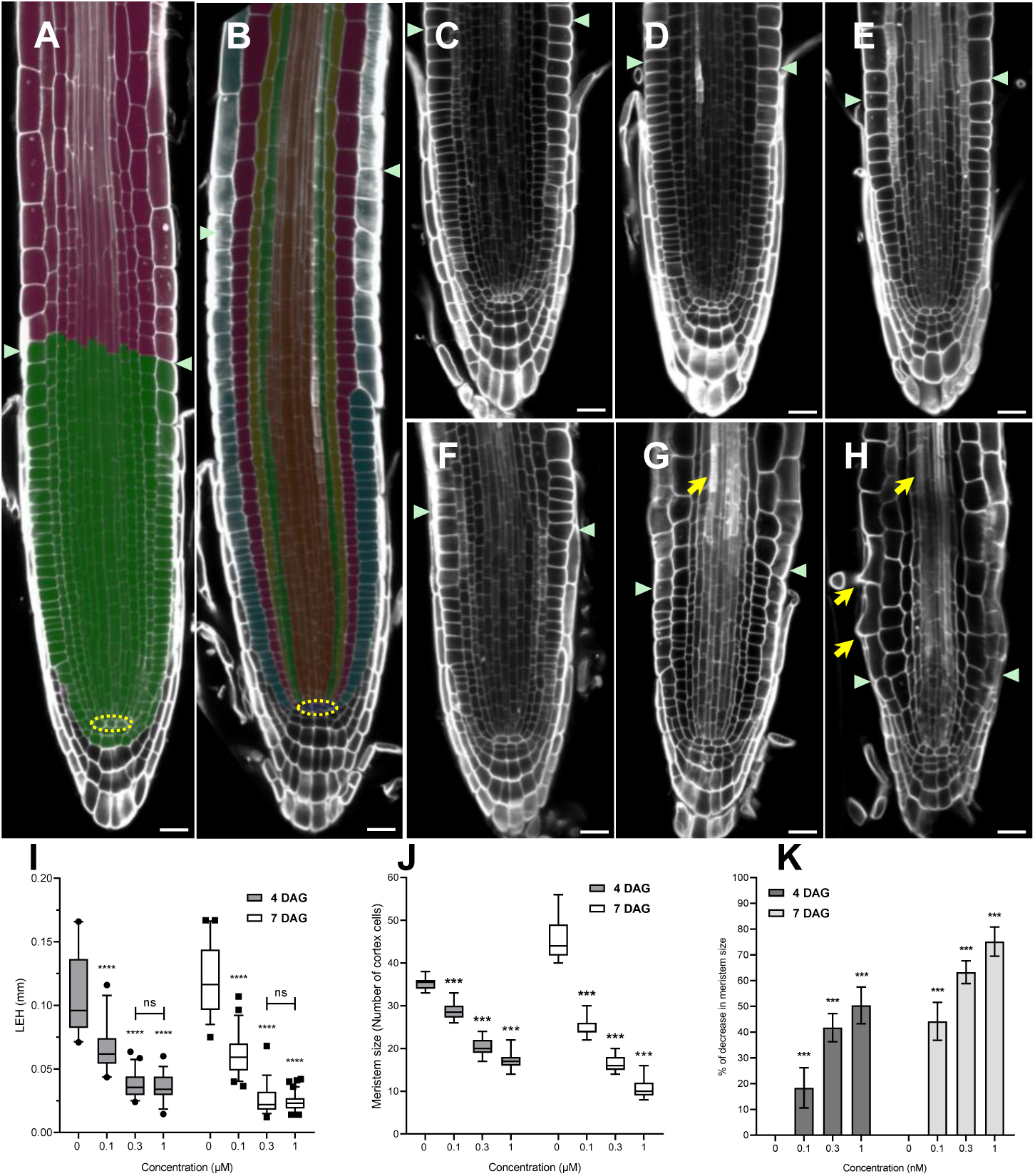
Lovastatin disturbs root developmental zones but not root radial organization. (A,B): Single root apex CLSM sections at 4 DAG and 7 DAG, respectively. Green arrowheads points to meristematic zone (green) boundary. Root meristem length is expressed as the number of cortex cells extending from the QC to the first elongated cortex cell (green arrowheads) (green meristematic zone, pink: elongation zone) (Perilli and Sabatini 2010) (A) and the histological patterning (from outer to inner layer: epidermis, cortex, endodermis, pericycle and stele). (C-E) Primary root patterning of seedlings 4 DAG treated with 0.1 μM (C), 0.3 μM (D) and 1 μM lovastatin (E). (F-H) Primary root patterning of 7 DAG seedlings treated with 0.1 μM (F), 0.3 μM (G) and 1 μM lovastatin (H). (I) Length of the first emerging root hair (LEH) at 4 DAG (0 μM N=15, 0.1 μM N=20, 0.3 μM N=20, 1 μΜ Ν=30) and 7 DAG (0 μM N=15, 0.1 μM N=20, 0.3 μM N=30, 1 μΜ Ν=30). Outliers represent values outside the 5-95%. Two-way ANOVA was used followed by Sidak’s post hoc analysis for multiple comparison. (J): Meristem size of 4 DAG (0 μM N=20, 0.1 μM N=20, 0.3 μM N=18, 1 μM N=19) and 7 DAG seedlings (0 μM N=14, 0.1 μM N=18, 0.3 μM N=15, 1 μM N=19). (K): Percentile % decrease of meristem size in comparison to untreated seedlings at 4 DAG (0 μM N=20), (0.1 μM N=20, 0.3 μM N=18, 1 μM N=19) and 7 DAG (0 μM N=14, 0.1 μM N=18, 0.3 μM N=15, 1 μM N=19). ***:*p*<0.001, ****: *p*<0.0001. Scale bars: 20 μm. N: number of seedlings. At least 3 biological replicates were used.

Four-day-old seedling roots showed normal root organization at all concentrations tested, although the lovastatin-treated roots were narrower than the mock (Fig 2A, C-E). After 7 d of lovastatin treatment, the histological pattern remained unaffected (Fig 2B, G-I), while cell number in each developmental zone decreased, possibly indicating a decline in mitotic activity (Fig. 2J). The final cell length was shorter than expected, as shown by the decreased length of the first emerging root hair (Fig 2I, Table 3), which probably contributed to overall decreased root length.

After 10 d of treatment, these effects were pronounced in roots treated with 0.1 μΜ-1μM lovastatin, while roots treated with 0.01-0.075 μM appeared almost unaffected (Supplementary Fig. S4). Exposure to higher concentration (0.3 μM and 1 μM) for 7 d resulted in shrinkage of the root apex developmental zones. This was indicated by the emergence of root hairs proximal to the root tip (Fig 2 G, H) and by early differentiation of protoxylem elements at *ca.* 180 μm from the tip (Fig 2 G, H arrows) in 1 μM lovastatin-treated seedlings.

Meristem size decreased in a dose-dependent and time-dependent manner across the lovastatin concentrations (Fig. 2J, K, Table 4, Table 5). After 4 d of treatment, 0.1 μM led to approximately 18% reduction of meristem size compared to the mock, 0.3 μM decreased root meristem size by an average 42%, whereas 1 μM reduced meristem size to half (Fig. 2K). The effect on meristem size was even more pronounced after 7 d of treatment, by which 0.1, 0.3 and 1 μM of lovastatin led to an average decrease of meristem size by 44, 63 and 75 %, respectively (Fig. 2K). The fact that the response to 0.1 μM was enhanced over time (average of 18% and 44% reduction of meristem size at 4 and 7 d of treatment, respectively) indicates that at low concentrations the effect accumulates with prolonged exposure, pointing towards a time-dependent effect. However, the response is more stable across the range of 0.1 to 1 μM. At both 4 and 7 d of treatment, the difference in meristem size between 0.1 and 0.3 μM is ∼21%, whereas between 0.3 and 1 μM meristem size reduced by ∼10% (Fig. 2K). This implies that the compound’s effect is not strongly time-dependent in that range; higher concentrations achieve near-maximal inhibition within the first 4 d and the system might be already reaching a plateau (Fig. 2J, K).

**Table 4.** The final cell length is significantly decreased. Length of the first emerging root hair (LEH) as determined by DIC images. Two-way ANOVA was used followed by Sidak’s post hoc analysis. N: number of seedlings. At least 3 biological replicates were analyzed.

|  | **0 μM** | **0.1 μM** | **0.3 μM** | **1 μM** |
| --- | --- | --- | --- | --- |
| **4 DAG** | 0.107±0.007<br>(N=15) | 0.06±0.003<br>(N=20) | 0.036±0.001<br>(N=20) | 0.035±0.001<br>(N=30) |
| **7 DAG** | 0.119±0.004<br>(N=15) | 0.06±0.002<br>(N=20) | 0.025±0.01<br>(N=30) | 0.023±0.0007<br>(N=30) |

**Table 5.**
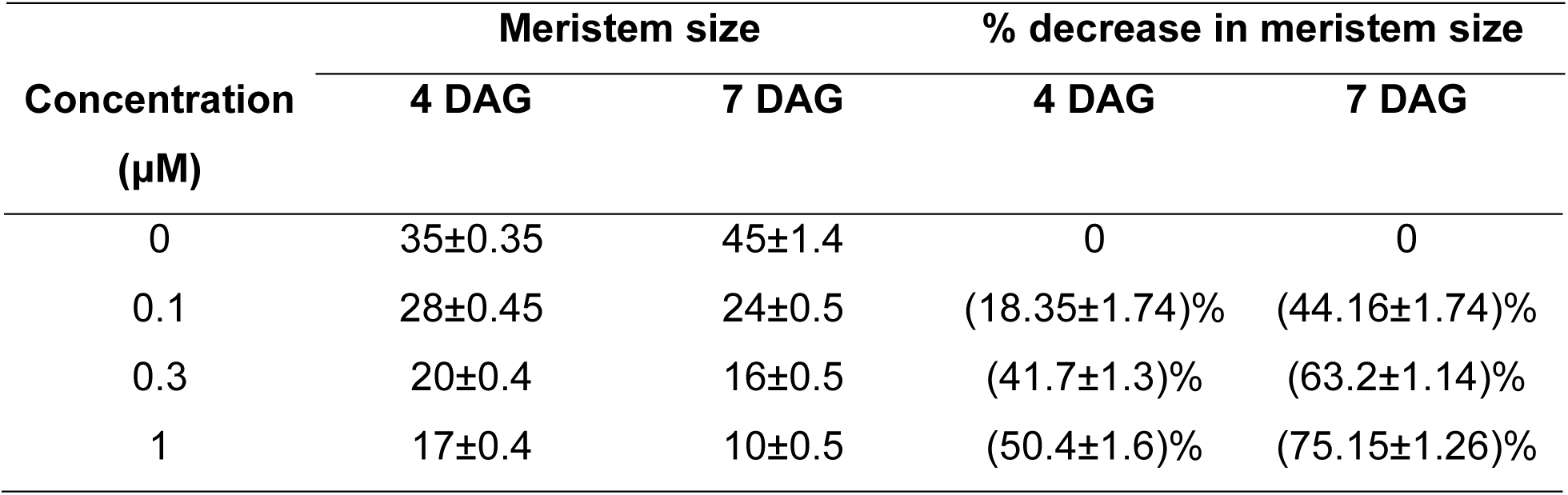
Meristematic size is highly affected by lovastatin treatment. Determination of meristem size according to Perilli and Sabatini (2010). Meristem size was determined from cleared and cell wall-stained seedlings 4DAG and 7DAG. % decrease of meristem size was analyzed compared to mock samples. Values represent mean±standard error.

These results indicate that the meristematic zone is highly sensitive to lovastatin. Even after four d at 1 μΜ the cells over the QC appeared elongated compared to the mock-treated cells (Fig. 2A, E). After 7 d in 0.3 μM and 1 μM lovastatin, the generation of new cells was almost completely inhibited, likely due to reduced cell division rate. Cells over the QC lost their typical meristematic identity and became elongated (Fig. 2B, G, H).

Despite inhibition of length, the radial organization of the primary root remained unaffected by lovastatin (Supplementary Fig. S5, Supplementary Fig. S6). Optical cross sections revealed no aberrations in the organization of the columella initials tier or the stele initials tier, or the QC cells (Supplementary Fig. S6). Since all cell layers were present in the cross sections at the beginning of the elongation zone (Supplementary Fig. S5A), the radial organization of the primary root was unaffected by lovastatin at any given concentration or treatment duration. Cell numbers within each cell layer did not exhibit any variation in all lovastatin concentrations at 4 DAG (Supplementary Fig. S6A), whereas at 7 d of treatment there was an unexpected decrease in atrichoblast number at 0.1 μM lovastatin and a decrease of stele cells at 1 μM lovastatin (Supplementary Fig. S6B).

Based on the severity of the observed effects, the 1 μM concentration was selected for all the following analyses. At this concentration all developmental and phenotypic abnormalities reached a plateau.

### Lovastatin disturbed lateral root primordium formation and lateral root emergence

To further investigate the effect of lovastatin on root development, lateral root emergence and growth were examined. Developmental stages of root primordia were identified in 10 d seedlings, mock and lovastatin-treated seedlings (Fig. 3). Lateral root primordia were classified accordingly (Malamy and Benfey, 1997). In lovastatin-treated seedlings, incomplete cell walls and ectopic positioning of several cells beginning at stage III were observed, ultimately leading to malformed emerging lateral root (Fig. 3 E, M). Interestingly, during the last stages of LRP development, extensive cell divisions occurred in the pericycle (Fig. 3 P, R-T), making the distinction and measurement of LRPs and lateral roots more tedious. Notably, when 4 DAG lovastatin-treated seedlings were transferred to mock medium, adventitious and lateral roots formed and elongated, compensating for the primary root growth impairment (Supplementary Fig. S3 I).

**Figure 3:**
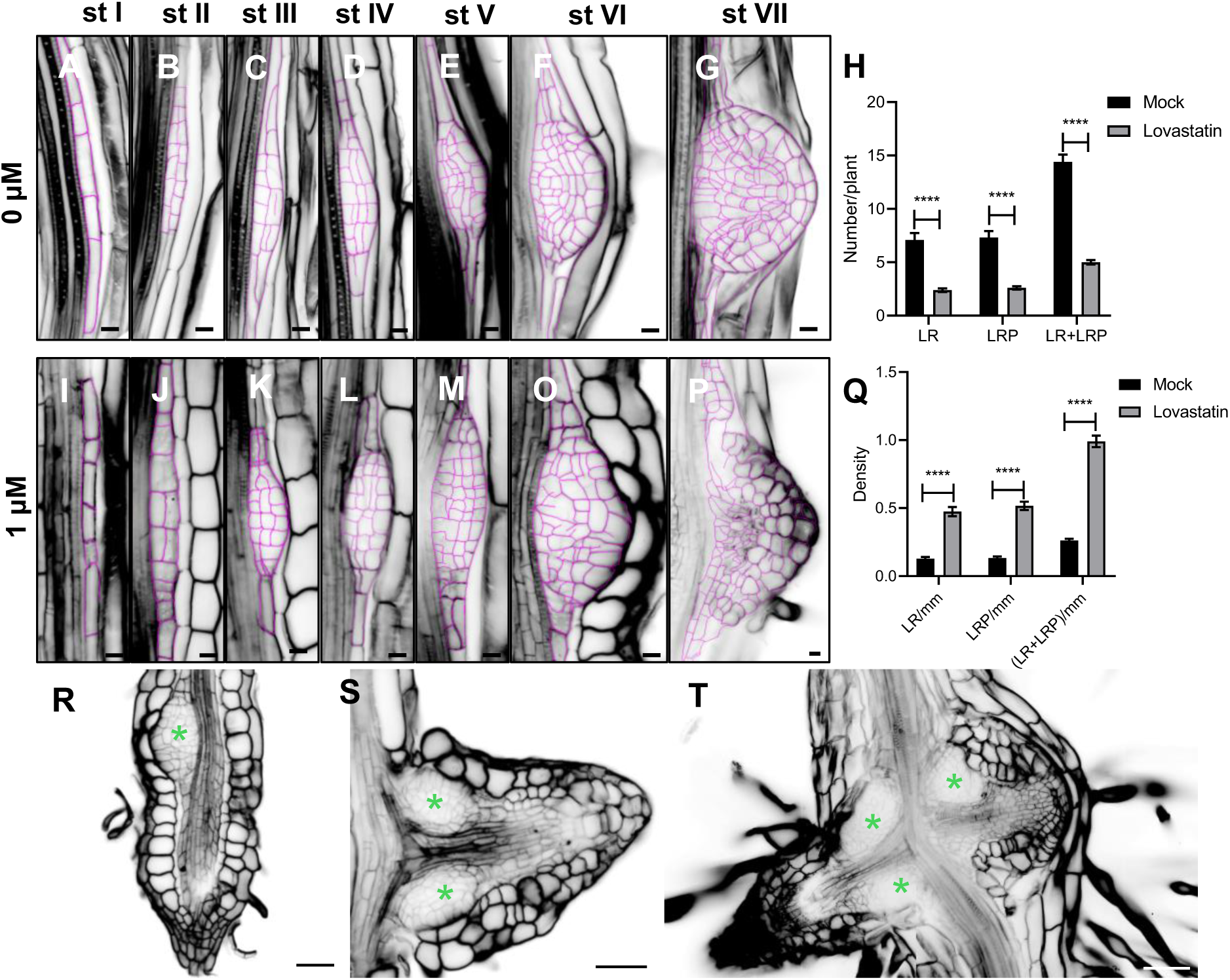
Disruption of lateral root primordium formation and lateral root emergence. (A-G): Calcofluor White-stained mock lateral root primordium developmental stage I-stage VII 10 DAG. (I-P): Lovastatin-treated lateral root primordium stage I-stage VII 10 DAG. (H): Lateral root primordia and lateral roots significantly decreased compared to untreated seedlings (N=22 for untreated, N=33 for 1 μM lovastatin-treated). (Q): Density of lateral root primordia significantly increased when treated with 1 μM (N=22 for untreated, N=33 for 1 μM lovastatin-treated). (R-S): Abnormal development of lateral root primordia, adjacent to primary root apex (R), either on underdeveloped lateral root (S) or formation of lateral root primordia on the adventitious roots as indicated by asterisks. **** p<0.0001. N: number or seedlings. For statistical analysis t-test was performed. At least 3 biological replicates were used.

Apart from LRP abnormalities, the number and density of LRPs and lateral roots were severely affected (Supplementary Fig. S7). The number of LRPs and lateral roots decreased (Fig. 3 H), while their density increased significantly (Fig. 3Q, Table 6). This observation indicates that the lovastatin-treated seedlings continue to respond to developmental cues related to lateral root formation despite the inhibition of sterol biosynthesis.

**Table 6:** Lateral root primordia (LRP) and lateral root (LR) quantification of untreated (mock) and treated (lovastatin) 10d old seedlings. Values represent mean±standard error. N: number of different seedlings used. Statistical significance was evaluated with two-way ANOVA.

|  | Mock | Lovastatin | Significance |
| --- | --- | --- | --- |
| LRP/plant | 7.3±0.61 (N=22) | 2.6±0.15 (N=33) | **** |
| LR/plant | 7.09±0.65 (N=22) | 2.39±0.16 (N=33) | **** |
| (LRP+LR)/plant | 14.4±0.68 (N=22) | 5±0.21 (N=33) | **** |
| LRP/mm | 0.13±0.01 (N=22) | 0.51±0.03 (N=33) | **** |
| LR/mm | 0.12±0.01 (N=22) | 0.47±0.03 (N=33) | **** |
| (LRP+LR)/mm | 0.26±0.01 (N=22) | 0.99±0.04 (N=33) | **** |

### Disturbance of auxin and cytokinin balance and distribution in lovastatin-treated seedlings

Auxin and cytokinin interaction coordinate various developmental cues in roots including patterning of the root and regulation of apical meristem (Schaller *et al*., 2015). Therefore, auxin and cytokinin distribution were examined in seedlings treated with lovastatin for 4 d and 7 d. Lovastatin-treated primary roots exhibited alterations in signal intensity and distribution of the auxin reporter line *DR5rev::GFP* compared to the mock (Fig. 4A, B). Specifically, lovastatin treatment resulted in depletion of *DR5rev::GFP* signal from the columella, while at the QC signal intensity was not affected in comparison with the mock (Fig. 4 A, B, E-G). In seedlings treated with lovastatin for 7 d, auxin distribution in both the root cap and the QC further reduced (Supplementary Fig. S8 A, B, F, G). Interestingly, the DR5rev-GFP signal increased in the stele, further supporting the severe disruption of auxin maxima gradient possibly affecting the LRP number and density (Supplementary Fig. S8 A, B, E; Supplementary Fig. S9).

**Figure 4:**
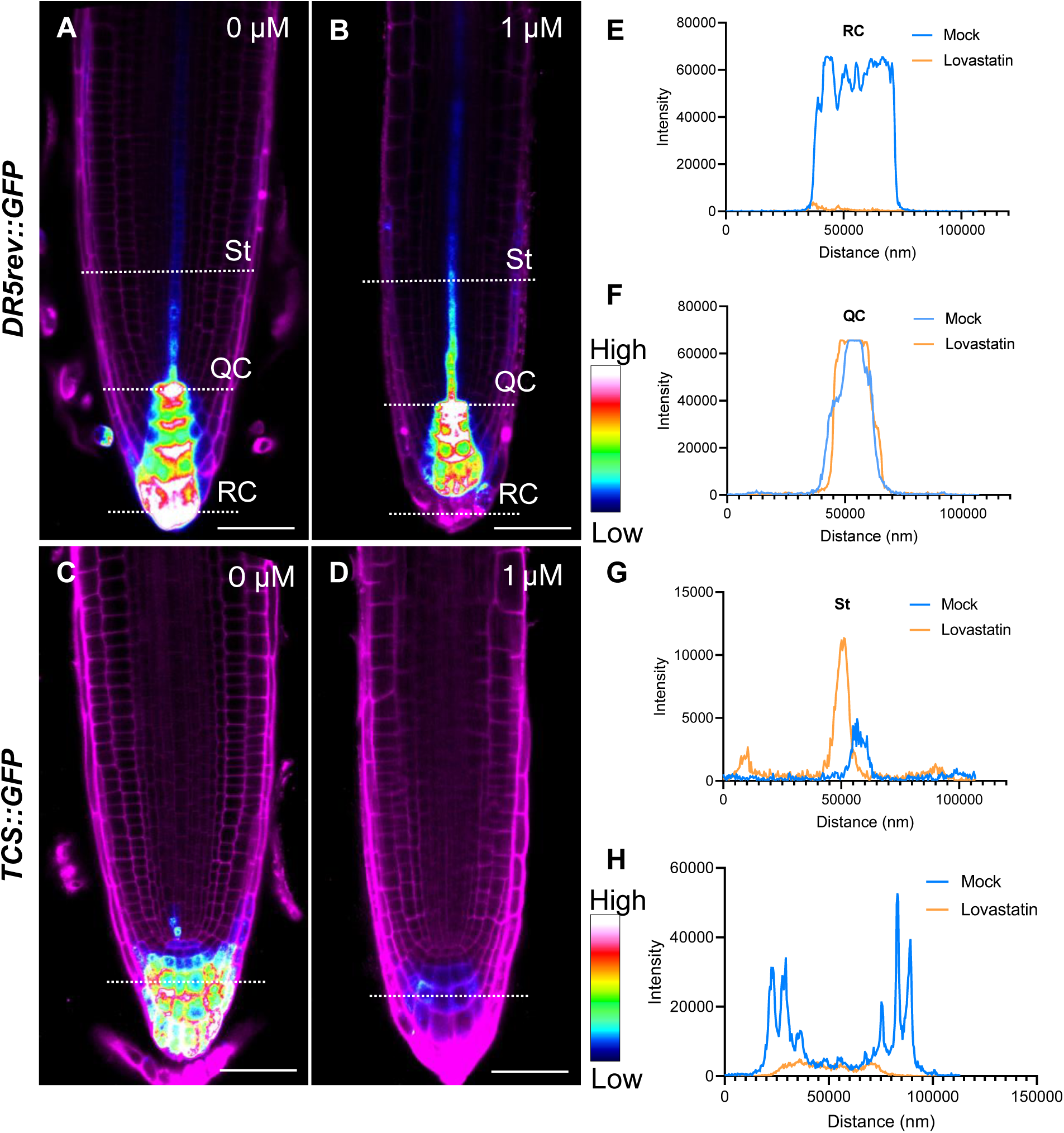
Lovastatin treatment disturbed auxin/cytokinin balance. Single CLSM sections of propidium iodide stained *DR5rev::GFP* root tips of 4 d untreated (A) and 1 μM lovastatin-treated seedlings (B). Propidium iodide stained *TCS::GFP* CLSM images of 4 d untreated (C) and treated (D) roots. Intensity profiles at the line positions of DR5rev::GFP on root cap (RC) (E), QC (F) and stele (St) (G) were altered after lovastatin treatment. Intensity profile of *TCS::GFP* was significantly decreased at the cytokinin collection spot (H). At least 10 seedlings were used for each condition. Scale bar: 50 μm.

Cytokinin signaling marker *TCS::GFP* was significantly decreased in root apices treated for 4 d with lovastatin, while after 7 d of exposure it almost completely abolished (Fig 4 C, D, H, Supplementary Fig. S8 C, D, H). This indicated either deficiency in cytokinin distribution or production.

### Lovastatin affects PIN expression and localization in primary root apex and lateral roots

The abnormal distribution of auxin suggested possible defects in auxin transport. To investigate this, the expression and localization patterns of major auxin efflux carrier proteins of the root, PIN1, PIN2, PIN3, PIN4 and PIN7 were analyzed. In roots treated for 4 d with 1 μM lovastatin PIN1 and PIN3 responsible for auxin efflux towards the root tip, showed increased gene and protein expression, compared to mock roots (Fig. 5A, B, C, Supplementary Fig. S10). Gene expression of PIN3 and PIN7 were significantly elevated while PIN2 and PIN4, remained unaffected (Fig. 5 A, Supplementary Fig. S10). Even though the total protein profile of lovastatin-treated *vs* untreated roots did not differ by silver staining (Supplementary Fig. S11), protein expression of PIN1 and PIN3 was significantly increased (*p*=0.0179 and *p*=0.0414 respectively, *n*=5), while PIN4 were unchanged (*p*=0.43) (Fig. 5 B, C).

**Figure 5:**
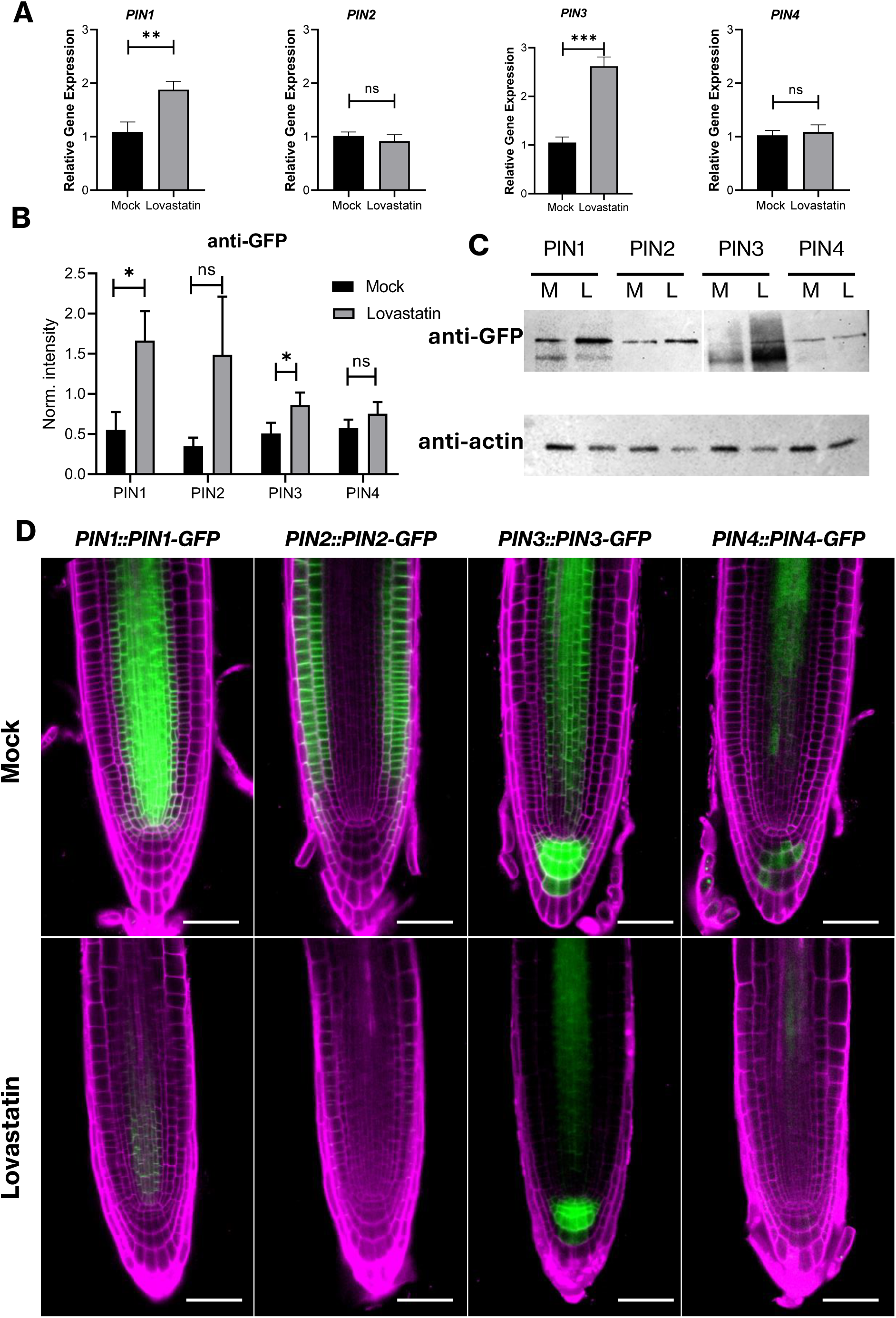
Lovastatin disturbs PINs localization and expression. (A) RT-qPCR analysis of PIN1, PIN2, PIN3 and PIN4 of mock (N=30) and 1 μM lovastatin-treated roots (N=90). (B) Semiquantitative analysis of blotted mock and treated roots of PIN1, PIN2, PIN3 and PIN4 GFP reporter lines. Equal loading was tested with actin. (C) Western blot analysis of 20 μg of total protein. (D) Single CLSM sections of propidium iodide-stained 4 d PIN reporter lines. Scale bars: 50 μm. At least 10 seedlings were used for each reporter line. *p<0.05, **p<0.01, ***p<0.001. T-test parametric test was used.

Notably, this increase in PIN expression contrasted with the reduced PIN signal observed by CLSM (Fig. 5 D). In mock seedlings, PIN1 localized to basal membranes of vascular cells (Fig. 5D, Supplementary Fig. S12 A), PIN2 to the apical membrane in the root cortex and epidermis (Fig. 5D, Supplementary Fig. S12 B), PIN3 to the first tier of columella cells distal to the columella initials at all faces of plasma membrane (Fig. 5D, Supplementary Fig. S12 C), and PIN4 showed weaker polar localization in the QC and distribution in stele (Fig. 5D, Supplementary Fig. S12 D).

Following lovastatin treatment, plasma membrane localization of all PINs was reduced as early as 4 DAG (Fig. 5D). By 7 days of treatment membrane localization was abolished except for a weak PIN3 signal in the stele (Supplementary Fig. S12). Together, the gene and protein expression data indicated increased PINs production while their reduced plasma membrane localization suggested protein trafficking issues or improper membrane targeting due to altered membrane fluidity. The increased expression might reflect a response to deficient localization and/or reduced degradation (Baster *et al*., 2013).

Next, since auxin efflux is essential for LPR development, functionally redundant *PIN1::PIN1-GFP* and *PIN3::PIN3-GFP* proteins were examined. At the first stage. PIN1 was localized on the anticlinal faces of the short initial cells (Supplementary Fig. S13 A-C). From stage II onward, PIN1-GFP signal was also found at periclinal faces and in later stages PIN1-GFP signal gradually increased towards the lateral root tip (Supplementary Fig. S13, Figure 6 A-D). In emerged lateral roots PIN1-GFP localized predominantly towards the root apex with a pattern similar to that of primary roots (Fig. 6 C, D). After lovastatin treatment, PIN1-GFP was almost absent from LRPs, yet present in primary root stele (Fig. 6 J-M). At emerged lateral roots, faint and diffuse PIN1-GFP signal was recorded in central LRP area with no membrane localization (Fig. 6 L, M).

**Figure 6:**
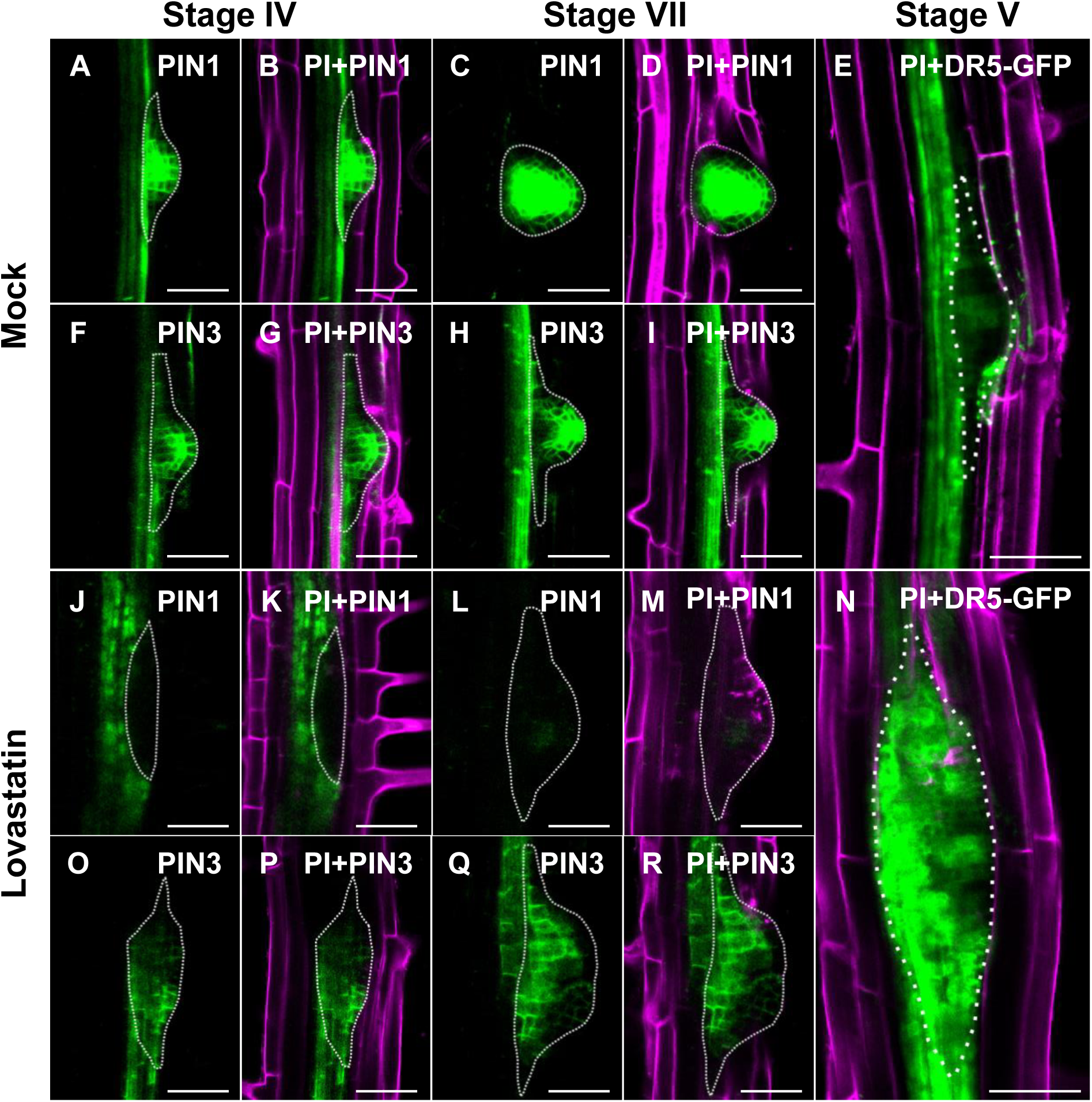
Lovastatin disrupts auxin gradient through disruption of plasma membrane localization of PIN1 and PIN3 during lateral root formation. (A-I) Untreated *PIN1::PIN1-GFP* (A, D), *35S::DR5-GFP*, (E) and *PIN3::PIN3-GFP* root primordia (F-I). (J-N) Effect of 1 μM lovastatin treatment to *PIN1::PIN1-GFP* (J, K), *35S::DR5-GFP* (N) and *PIN3::PIN3-GFP* (O-R). Plasma membrane localization of PIN1-GFP is completely abolished at stage IV with signal preserved in stele (A,B compared to J, K). At stage VII PIN1-GFP has low expression and no membrane localization (C, D compared to L, M). (E, N) Increased DR5-GFP signal at the primordium initials at stage V in treated seedlings. PIN3-GFP gradually establishes polarized localization towards developing lateral root (F-I) while in lovastatin treated seedlings reduced plasma membrane signal is observed without any specific polarization (O-R). Seven-day old seedlings were used. Dotted line mark the lateral root primordium. PI: propidium iodide. Scale bar: 50 μm.

In mock seedlings, PIN3-GFP at the early stages of LRP development was localized at the anticlinal faces of short initial cells and from stage II at the periclinal faces (Supplementary Fig. S13 D-F). From stage IV, PIN3-GFP expression localized towards the columella precursors of the newly forming meristem (Fig. 6 F-I). After lovastatin treatment PIN3-GFP signal remained on plasma membrane but its polarity was disrupted (Fig. 6 O-R). Extensive cell divisions in the pericycle, similar to those observed in *pin* mutants (Benková *et al*., 2003), made it more difficult to recognize the different developmental stages of LRPs (Fig. 6 O-R). The abnormal PIN1-GFP and PIN3-GFP expression coincided with the disrupted auxin accumulation (Fig. 6 E, N). Normally, *DR5rev::GFP* is active at all stages of LRPs (Fig. 6 E). However, in lovastatin-treated seedlings, DR5rev showed increased diffusion without clear separation of LRP formation (Fig. 6 N). The increased auxin signaling together with defects in PIN polarity demonstrated that PIN-dependent auxin efflux possibly affected the abnormal LRP formation.

### Effect of lovastatin on PIN2 dynamics and lateral diffusion

PIN auxin transporters generate auxin gradients that drive primary root development and lateral root emergence. In particular, PIN2 acts at epidermal (with shootward polar localization) and cortical root cells (with a rootward polar localization) to regulate proper lateral root spacing. As lovastatin causes ectopic lateral root formation and abortive initiation, also considering the membrane deficiencies and the discrepancy between PIN2-GFP localization and abundance after lovastatin treatment, PIN2 mobility in root cells was examined with FRAP analysis.

For this purpose, seedlings stably expressing *PIN2::PIN2-GFP* were grown on solid medium containing 1 μM lovastatin for 4 d and subjected to FRAP analyses. PIN2-GFP showed slow mobility in mock samples (mobile fraction: 22.48±7.23%, immobile fraction: 77.52±7.23%, recovery rate t1/2: 13.62± 5.26 s, Fig. 7A-E, Supplementary Vid. S1). FRAP of lovastatin-treated seedlings demonstrated an alteration of membrane dynamics. The mobile fraction was significantly decreased (mobile fraction: 16.1±4.3%, immobile fraction: 83.9±4.3%, *p*=0.016 Welch’s test) indicating that the proportion of the molecules that are free to move is lower, compared to mock samples (Fig. 7 A-E). The PIN2-GFP recovery rate of lovastatin treated seedlings was significantly lower (t1/2: 7.3±7.27 s *p*<0.0001), indicating membrane fluidity issues possibly due to disruption of sterol biosynthesis pathway and the presence of structural sterols in the plasma membrane (Fig. 7 E, Supplementary Vid. S2). FRAP analyses showed that PIN2-GFP has faster recovery rate but subsequent decrease in lateral mobility in the presence of newly adapted PIN2-GFP molecules to the plasma membrane (Fig. 7 C, E). This might indicate alterations in plasma membrane structural composition and is in accordance with the PIN localization experiments.

**Figure 7:**
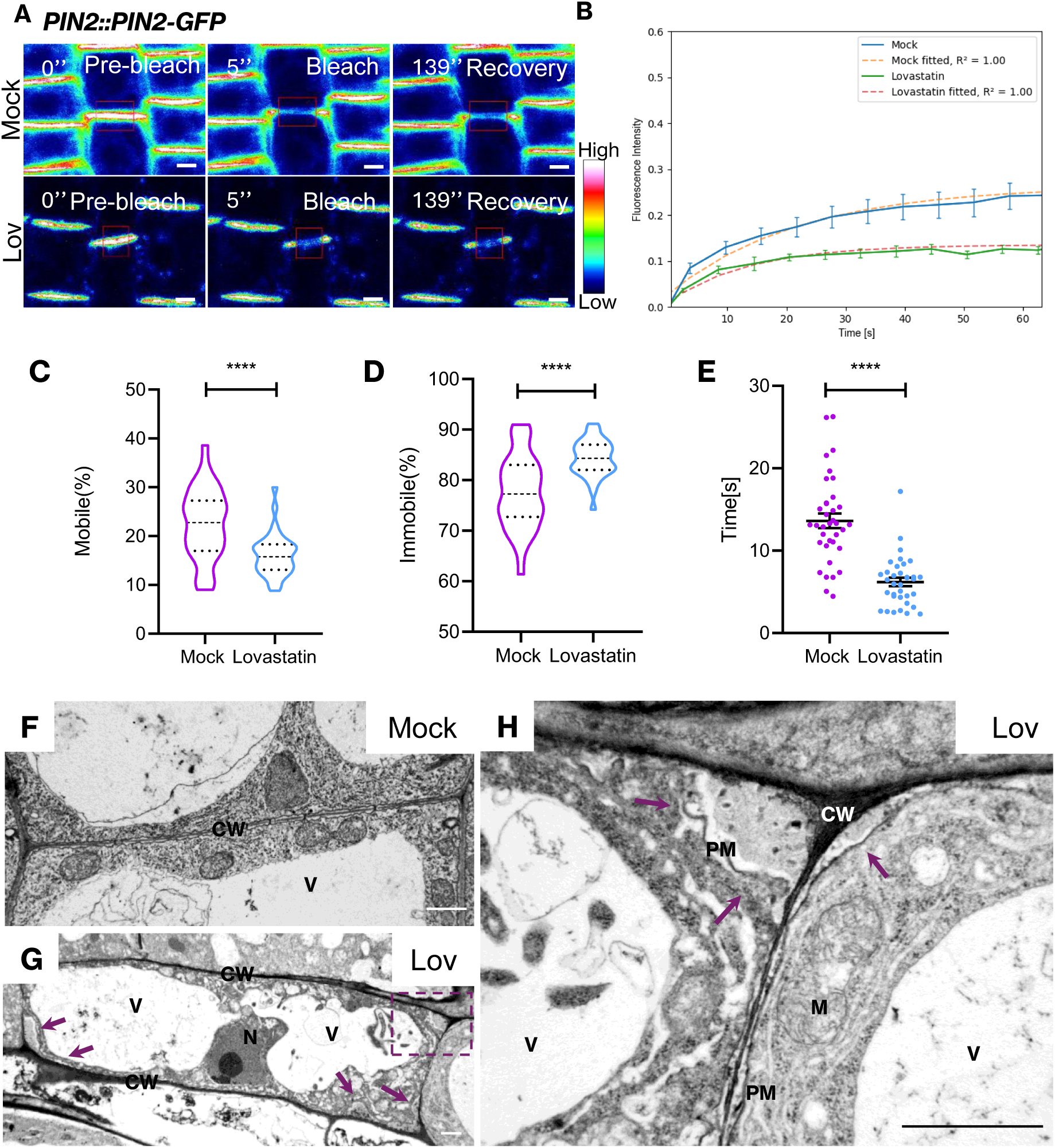
Alteration of PIN2 dynamics and lateral diffusion on 4 d and 7 d lovastatin-treated seedlings and disruption cell wall-PM continuum. (A): Representative FRAP experiment of root plasma membrane of seedlings untreated (N=35) or 1 μM lovastatin-treated (N=35) for 4 d. Red rectangle represent bleached region of interest. Pre-bleached, bleached and timepoint (in sec) during recovery are shown. (B): Quantitative analysis of normalized FRAP. Data points and error bars represent mean±standard deviation from at least 3 different representative set of regions. Dashed lines represent fitting curve. Pre-bleach intensities were set to 1 and immediate post-bleach to 0. (C, D): Quantitative FRAP analysis of mobile (C), immobile (D). Dashed line correspond to median and dotted lines the quartiles. (E): Half time recovery rate (t1/2). Time point distribution and mean±standard error are presented (F-H): TEM micrographs of cortex cells of seven-day-old seedlings germinated and grown in mock medium (F) or 1 μM lovastatin (G, H). (H) Higher magnification of rectangle selected region of (G). Plasma membrane exhibits local detachment from the cell wall and deposition of amorphous cell wall components (arrows) (G, H). CW: cell wall, V: vacuole, N: nucleus, M: mitochondrion. Scale bar: 5 μm (A) and 1 μm (F-H). **** p>0.0001. t-test was used.

### Lovastatin compromised cell structural integrity

The defects in PIN2 localization and mobility, previously recorded here, imply that lovastatin treatment possibly also results in membrane alterations at the ultrastructural level. Therefore, root cell ultrastructure was examined by TEM.

After 5 d of treatment, the plasma membrane appeared detached from the cell wall at several sites (Fig. 8 A, B). After 7 d of treatment, plasma membrane detachment was further expanded, while the emerging gap was filled with amorphous cell wall material (Fig. 7 F-H, Fig. 8 C). Such pockets appear in different cell types, including epidermal and cortical cells and occur at membrane areas adjacent to either cross- or longitudinal cell walls without any specific prevalence (Fig. 7 F-H). Apart from the above, incomplete cell walls, probably resulting from hampered vesicle cohesion during cytokinesis, were occasionally observed (Fig. 8 D). Interestingly, in several root cells the nuclear envelope appeared dilated, exhibiting broader than normal perinuclear space, which in some cases was extremely swollen (Fig. 8 E, F).

**Fig. 8:**
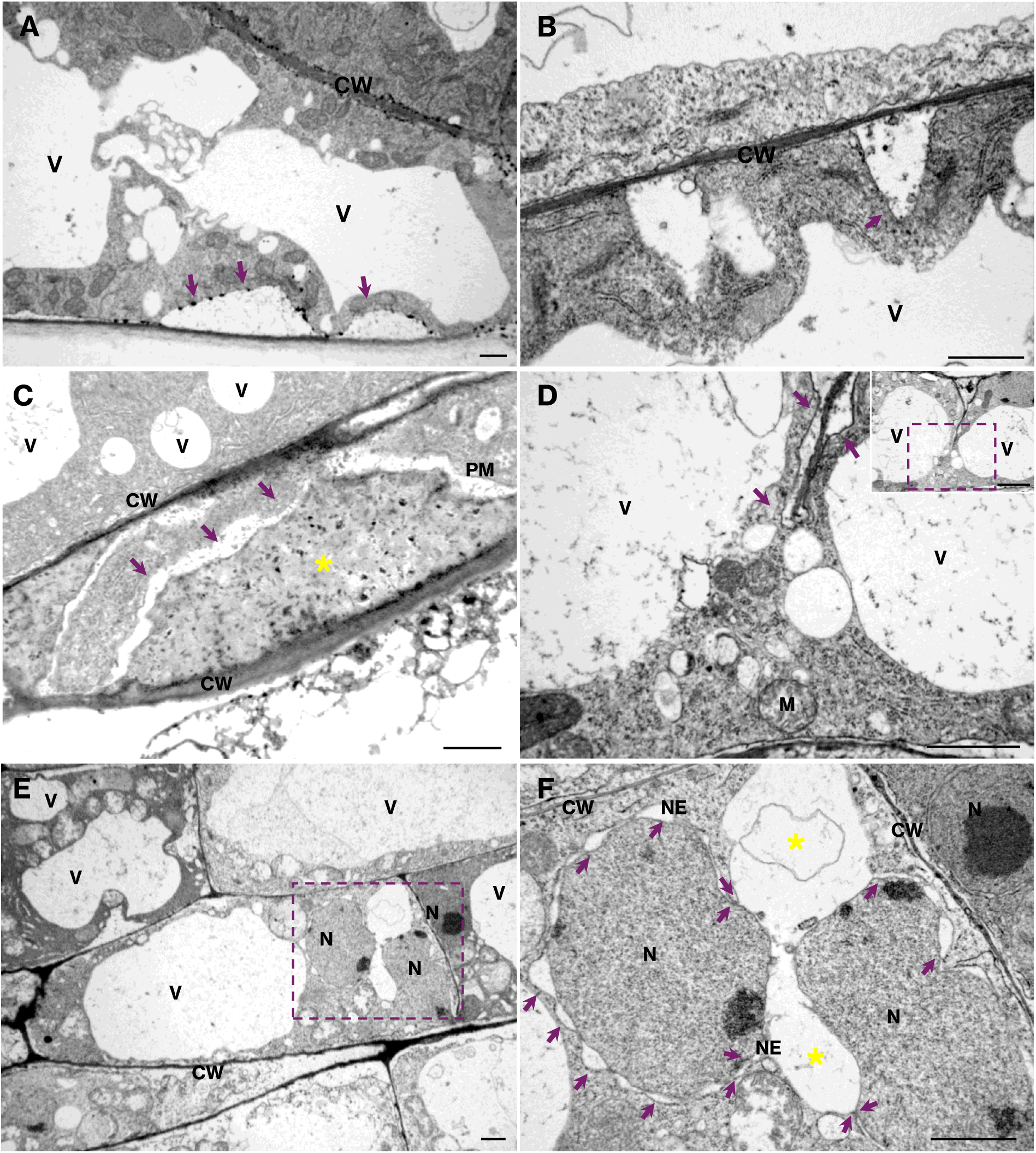
Lovastatin treatment causes local plasma membrane detachment, aberrant cell wall deposition and dilation of nuclear envelope in root cells. (A, B): TEM micrographs of root cells of five-day-old seedlings germinated and grown in 1 μM lovastatin. Plasma membrane exhibits local detachment (arrows) from the cell wall. (C-F) Prolonged (seven day) lovastatin treatment. (C): Extensive plasma membrane detachment (arrows), the area of which is full of amorphous cell wall material (yellow asterisk). (D): Incomplete cell wall with plasma membrane detachment sites (arrows) within the cytoplasm of interphase cell (the cell is shown at low magnification in the inset). (E): Extreme nuclear envelope dilation can be observed in this root cell (area included in rectangle). (F): Higher magnification of the above area. The arrows point to dilation sites and the asterisks at site of extreme perinuclear space swelling. CW: cell wall, V: vacuole, N: nucleus, NE: nuclear envelope, PM: plasma membrane, M: mitochondrion. Scale bar: 1 μm.

## Discussion

Here, the impact of sterol depletion on root system architecture by pharmacological inhibition of the mevalonate pathway was investigated. Prolonged lovastatin treatment at moderate concentrations severely compromised vegetative growth of *A. thaliana* seedlings, reflecting aberrations in cell ultrastructure and severe defects in primary root development and lateral root formation. Following a latent period of normal seedling growth, during which developing seedlings possibly consume sterols from seed resources (Doner *et al*., 2025). Thereon, lovastatin causes a significant retardation of primary root elongation, culminating to a stunted root. Tissue patterning was not affected nor alterations in cell division plane orientation were observed. However, basipetal growth of the root was disturbed, resulting in reduction of meristem size and occurrence of significantly short bulging trichoblasts close to the root tip. In parallel, lovastatin treatment perturbed root branching, resulting in ectopic lateral roots and abortive LRPs, largely inhibiting lateral root formation. Accordingly, the distribution and gradient of auxin and cytokinin were considerably affected and localization patterns of PIN auxin transporters were altered both in primary and lateral roots. Although the abundance of PINs was variably if at all affected, membrane kinetics of PIN2 through FRAP analysis showed decreased recovery rates after lovastatin treatment.

### Sterol depletion restricts meristem size and hampers primary root elongation

The stunted growth of both primary root and shoot is a dominant effect of lovastatin in *A. thaliana* and other plants (Nagata *et al*., 2002; Rodríguez-Concepción *et al*., 2004; Suzuki *et al*., 2004; Wang *et al*., 2023). Importantly, this effect is reversible upon removal of lovastatin, indicating that sterol depletion compromises developmental progression without causing irreversible damage or lethality. These findings are consistent with the well-established effect of lovastatin in reducing total sterol abundance without altering sterol composition (Suzuki *et al*., 2004; Kobayashi *et al*., 2007; Wang *et al*., 2023).

Lovastatin is a statin inhibiting 3-hydroxy-3-methylglutaryl coenzyme A (HMG-CoA) reductase (HMGR), acting early in the sterol synthesis pathway. In *A. thaliana* both HMGR1 (AT1G76490) and HMGR2 (AT2G17370) are highly expressed in late meristem, transition and elongation zones, whereas expression is low in the QC (Mironova, 2022). This spatial expression pattern provides a plausible explanation for the observed sensitivity of elongation-associated tissues to lovastatin, while meristem patterning itself remains largely preserved, likely supported by residual seed-derived sterols sufficient for QC establishment (Doner *et al*., 2025).

Higher lovastatin concentrations (≥0.3 μM) induced loss of meristematic cell identity above the QC, in epidermal and cortical tissues, consistent with premature differentiation. The reduction in meristem and elongation zone length and the appearance of shorter bulging trichoblasts closer to root apex suggests that lovastatin restricts both cell division and cell elongation. Comparable cell cycle arrest effects have been reported in plant and animal systems following lovastatin treatment, including arrest in G1 phase and reduced mitotic activity in early sterol biosynthesis mutants (Jakóbisiak *et al*., 1991; Rao *et al*., 1999; Souter *et al*., 2002; Hartig and Beck, 2005; JavanMoghadam-Kamrani and Keyomarsi, 2008; Mialoundama *et al*., 2014; Wang *et al*., 2023).

### Sterol content modulates auxin-cytokinin balance in root apex

The reduction in meristematic activity prompted the examination of hormone signaling. Lovastatin treatment caused pronounced alterations in auxin gradient and cytokinin abundance as revealed by *DR5rev::GFP* and *TCS::GFP.* Auxin acts through the development of gradients and concentration maxima providing the positional cue for developmental processes including primordia formation (Benková *et al*., 2003; Benjamins and Scheres, 2008).

Shift in auxin maxima in the QC and increased DR5rev-GFP signal along stele, as observed using *DR5rev::GFP,* align with previous published work in early steps of sterol biosynthesis mutants, including *hyd2/fk*, *smt1*, *smo1-1smo1-2* and *smo2-1smo2-2* mutants. In these lines, altered auxin signaling was associated with the reduced and/or modified sterol composition (Souter *et al*., 2002; Willemsen *et al*., 2003; Zhang *et al*., 2016; Song *et al*., 2019). Furthermore, other studies have suggested an interdependence between plasma membrane sterols and auxin signaling (Pan *et al*., 2009; Zhang *et al*., 2016).

The reduction of *TCS::GFP* may reflect crosstalk between cytokinin biosynthesis and the mevalonate pathway, potentially impacting cytokinin-related processes (Rodríguez-Concepción *et al*., 2004). Cytokinins are generally known to promote root apical meristem differentiation while inhibiting mitotic cell divisions (Schaller *et al*., 2014). Root architecture defects in the presence of lovastatin, did not display characteristics typically associated with cytokinin depletion, further suggesting that these are not predominantly driven by cytokinin depletion (Werner *et al*., 2003).

Although manipulation of brassinosteroids affects the localization patterns of TCS (Ohashi-Ito *et al*., 2023) there is no indication of brassinosteroid involvement herein. Possibly, disturbances in TCS-GFP localization following lovastatin treatment are caused by reduced structural sterols (Nakamoto *et al*., 2015; Zhang *et al*., 2016; Vogel *et al*., 2025). Interference with brassinosteroids production and signaling seems to exert opposite effects in overall root development, different than those observed after lovastatin treatment or in the HMGR1 mutant *hmg1* (Suzuki *et al*., 2004; Al-Mamun *et al*., 2024).

### Lateral root defects arise from mis-regulation of auxin gradients and PIN localization

Lovastatin also affected root branching by inhibiting lateral root formation. Notably, many LRPs formed but, failed to progress beyond early developmental stages and frequently arose at ectopic positions along the primary root or at the base of emerging lateral roots. These abortive LRPs likely correspond to previously described tumor-like structures observed in lovastatin-treated roots (Kobayashi *et al*., 2007).

Mis-regulation of lateral root formation is likely related to altered auxin distribution. Lovastatin treatment induced auxin accumulation in the stele, potentially providing inappropriate initiation signals for LRP formation (Benková *et al*., 2003). However, successful LRP development and emergence require tightly regulated auxin efflux by PIN transporters, especially PIN1 and PIN3 (Benková *et al*., 2003; Adamowski and Friml, 2015; Vilches-Barro and Maizel, 2015; Omelyanchuk *et al*., 2016). Accordingly, lovastatin treatment compromised the plasma membrane localization and polarity of PIN1 and PIN3, thereby likely impairing directional auxin transport, essential for LRP patterning and emergence. Similar defects in PIN localization and auxin distribution have been reported in sterol biosynthesis mutants (Souter *et al*., 2002; Willemsen *et al*., 2003), further supporting a functional link between sterol homeostasis and auxin transport (Kobayashi *et al*., 2007; Wang *et al*., 2023).

Mutations in sterol biosynthesis genes upstream of the 24-ethyl/24-methyl sterol branch point are known to disrupt PIN localization and auxin distribution, possibly affecting lateral root formation (Souter *et al*., 2002; Willemsen *et al*., 2003; Zhang *et al*., 2016; Boutté and Jaillais, 2020). In this context, the restoration of lateral root emergence in recovery from lovastatin treatment experiments, is noteworthy. A similar phenomenon has been described in the *smt2smt3* sterol biosynthesis mutant, where compromised lateral root development and auxin distribution was partially rescued by exogenous supplementation with β-sitosterol (Nakamoto *et al*., 2015). Together these observations are supportive of a role of structural sterols in maintaining auxin-dependent processes, rather than acting solely as static membrane components, a concept further reinforced by other reports (Zhang *et al*., 2016).

### Sterols regulate PIN dynamics through membrane integrity and trafficking

A central finding of this study is the comprehensive mis-localization of all analyzed PIN (PIN1, PIN2, PIN3 and PIN4) upon sterol inhibition. FRAP analysis revealed reduced membrane recovery rates for PIN2, indicating impaired membrane dynamics. In addition, lovastatin treatment caused partial plasma membrane detachment from the cell wall and defects in cell wall integrity, phenomena previously reported in sterol biosynthesis mutants (Schrick *et al*., 2004).

Membrane detachment may be related to PIN2 targeting issues and kinetic behavior, although this has to be proved. Other studies have also demonstrated that membrane fluidity reduction is a general property following genetic or pharmacological sterol depletion, not limited to PIN proteins but also including other membrane species as well (Willemsen *et al*., 2003; Men *et al*., 2008; Yang *et al*., 2013).

The mechanism by which sterols influence protein localization remains a subject of debate. While Zhang et al. (2016) argued that sterol composition, rather than quantity, is critical for PIN1 stability, present findings align with the “trafficking” hypothesis. Yang et al. (2013) demonstrated that sterols are essential for the trafficking of ABCB19 from the trans-Golgi to the PM. Alternatively, the “lipid raft” hypothesis proposed by Willemsen et al. (2003) suggests that sterols are required for the tethering of proteins into specific membrane microdomains. These data suggest that lovastatin-induced sterol depletion likely affects both: disrupting the initial trafficking of transporters PINs to the membrane, while simultaneously destabilizing those already present by altering membrane fluidity or microdomain integrity.

### Conclusion

In conclusion, the present study provides comprehensive evidence that sterol homeostasis is a fundamental requirement for the spatial regulation of auxin transporters and, consequently, proper plant development. By utilizing lovastatin to inhibit sterol biosynthesis, a disruption of auxin-cytokinin balance and global disruption of PIN protein localization was demonstrated, which correlates with a reduction of mitotic activity and compaction of developmental zones. These findings reconcile several disparate observations in the literature by suggesting that the “trafficking” and “microdomain” hypotheses are not mutually exclusive. Instead, a model is proposed where sterols act as master regulators of the plasma membrane environment-facilitating both the initial delivery of auxin transporters from the trans-Golgi network and their subsequent stabilization within lipid rafts. Ultimately, this work underscores the intricate synergy between membrane lipid composition and hormone signaling. Future research should focus on identifying specific sterol-protein interactions that govern these docking mechanisms, potentially through high-resolution lipidomic and proteomic profiling of membrane microdomains. These insights will be crucial for understanding how plants modulate their development in response to environmental and metabolic cues.

## Supporting information

Supplementary Vid S1

Supplementary Vid S2

## Supplementary data

The following supplementary data are available at JXB online.

**Supplementary Table S1**. Reporter lines used in this study.

**Supplementary Table S2.** Primers used in this study.

**Supplementary Figure S1: Screening of effect different lovastatin concentrations to seedling and root growth and rate.**

**Supplementary Figure S2: Screening of effect different lovastatin concentrations to shoot and hypocotyl.**

**Supplementary Fig. S3. Lovastatin treated seedlings recover after transplanting to control medium**.

**Supplementary Fig. S4: Lovastatin affects root developmental zones but not the histological patterning in a dose-dependent manner**.

**Supplementary Fig. S5. Radial organization of primary roots is not affected by any lovastatin treatment applied.**

**Supplementary Fig. S6: Lovastatin did not affect total cell number across the root cell layers except atrichoblasts and stele at 7 DAG.**

**Supplementary Figure S7. Inhibition of lateral root emergence after lovastatin treatment.**

**Supplementary Fig. S8. Lovastatin treatment disturbed auxin gradient and almost completely abolished cytokinin at 7 DAG.**

**Supplementary Fig. S9: Auxin gradient is significantly disrupted after lovastatin treatment.**

**Supplementary Fig S10. PIN7 increased gene expression after lovastatin treatment.**

**Supplementary Figure S11. Total protein cell lysate profile of PINs 4DAG of untreated (M) and treated with 1 μM lovastatin (L).**

**Supplementary Figure S12. PINs membrane localization is severely compromised at 7 DAG.**

**Supplementary Fig. S13. Expression of PIN1-GFP and PIN3-GFP in early stages of lateral root development in mock seedlings.**

**Supplementary Video S1. Representative FRAP of mock *PIN2::PIN2-GFP* 4DAG**

**Supplementary Video S2. Representative FRAP of 1 μM lovastatin treated *PIN2::PIN2-GFP* 4DAG.**

## Acknowledgements

We thank Nottingham Arabidopsis Stock Centre for the reporter lines.

## Author contributions

Conceptualization (VG, GK), data curation (VG, CT, SP), formal analysis (VG, CT, GK), methodology (VG, CT, AK, SP, AA, EP, GK), software (AA), resources (GK, KV), supervision (GK), funding (KV, EP, GK), writing (VG, EP, GK).

## Data availability

All data supporting these findings are available in this paper and the supplementary material published online. FRAPedia releases and source code are available on GitHub: https://github.com/AlkiviadisAthanasiadis/FRAPedia.git.

## Abbreviations

ANOVA: analysis of variance
DAG: days after germination
DIC: differential interference contrast microscopy
HMGR: 3-hydroxy-3-methylglutarylCoA reductase
FRAP: fluorescence recovery after photobleaching
LRP: lateral root primordium
NA: numerical aperture
QC: quiescent centre
RC: root cap
St: stele
SD: standard deviation

