## SupplementaryFigures for "Effects of lovastatin on auxin transport and root development in *Arabidopsis thaliana*"

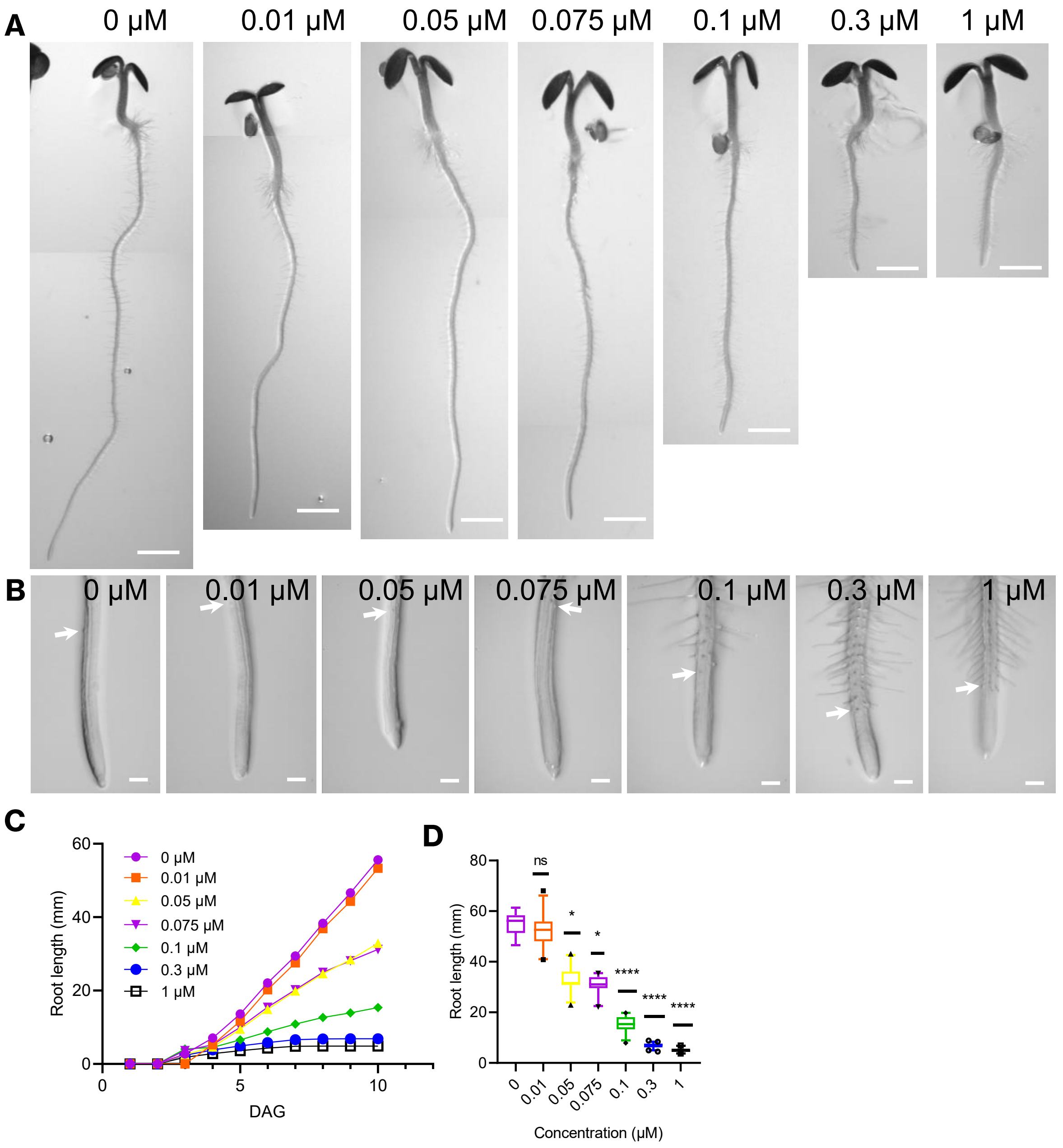


**Supplementary Figure S1: Screening of the effect of increasing lovastatin concentrations to seedling and root growth and rate.** (A) Five-day-old seedlings grown in increasing concentrations of lovastatin. (B) Respective root apices from (A) at higher magnification. Arrows point to the first bulging root hair. (C) Root growth rate at various lovastatin concentrations for 10 d. 0 μM: N=18, 0.01 μM N=29, 0.05 μM: N=26, 0.075 μM: N=23, 0.1 μM: N=27, 0.3 μM: N=22, 1 μM: N=43. (D) Root length of seven-day-old seedlings grown in various lovastatin concentration. Kruskal's Wallis test combined with Dunn's multiple comparison. For 0 μM N=17, 0.01 μM N=17, 0.05 μM N=23, 0.075 μM N=22, 0.1 μM: N=26, 0.3 μM: N=42, 1 μM: N=21. Comparisons were made against 0 μM (mock). Bars represent 5-95% of sample distribution and outliers are shown. * $p$<0.05, **** $p$<0.0001 Scale bar: 1 mm (A), 0.1 mm (B). N: number of different plants. At 3 biological replicates were used.

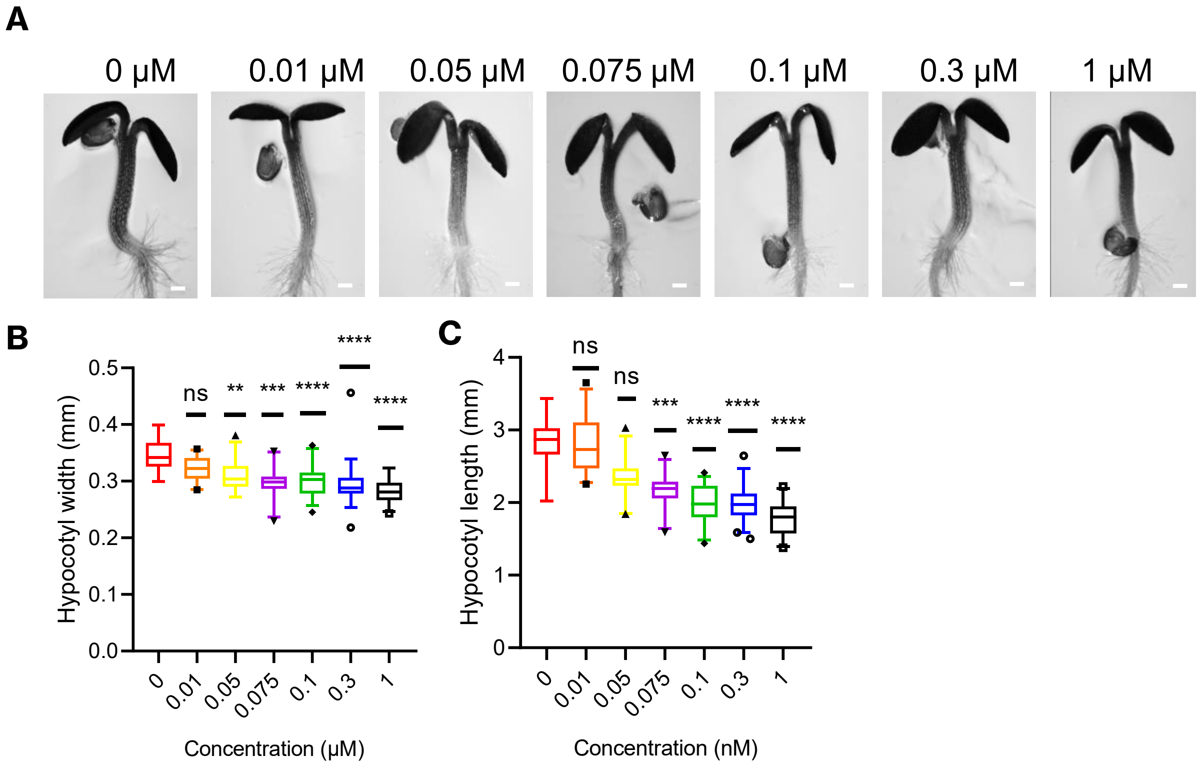


**Supplementary Figure S2: Screening of the effect of increasing lovastatin concentrations to the shoot and hypocotyl.** (A) Shoots of five-day-old seedlings grown in increasing lovastatin concentrations. (B) Hypocotyl length is significantly reduced in seedlings treated with lovastatin ≥0.05 µM. (C) Hypocotyl width is affected by lovastatin treatment significantly at concentration higher than 0.01 µM. For (B) and (C): 0 µM N=17, 0.01 µM N=17, 0.05 µM N=23, 0.075 µM N=22, 0.1 µM: N=26, 0.3 µM: N=42, 1 µM: N=21. Bar plots represent 5-95% of sample distribution and outliers are shown. * $p<0.05$, ** $p<0.01$, *** $p<0.001$, **** $p<0.0001$ Scale bar: 0.2 mm. N: number of different plants. At 3 biological replicates were used.

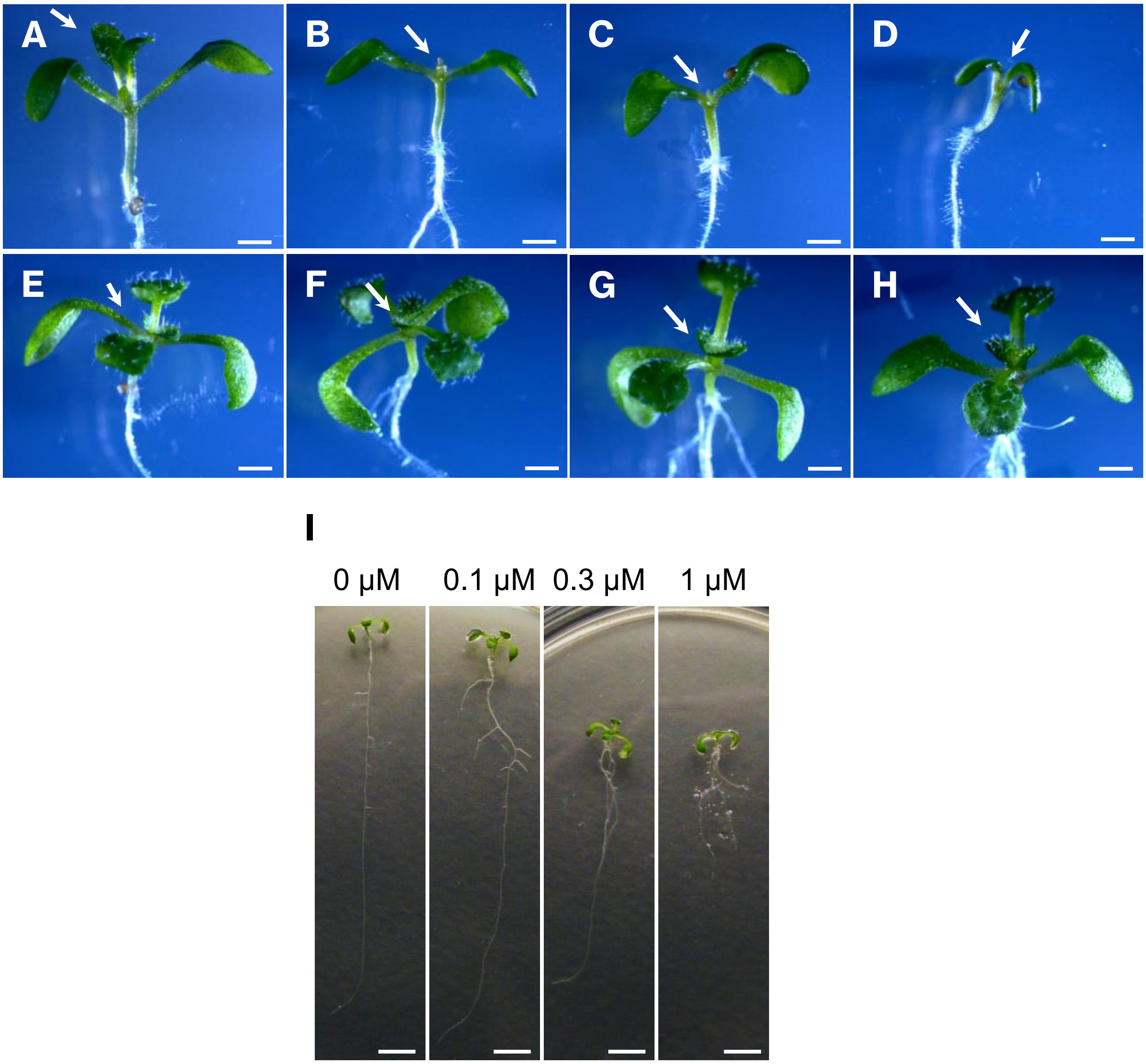


**Supplementary Fig. S3. Lovastatin-treated seedlings recover after transplanting to control medium**. (A-D) Shoot apical meristem (arrow) of ten-day-old seedlings is repressed by lovastatin treatment [(A) 0 μM, (B) 0.1 μM, (C) 0.3 μM, (D) 1 μM]. (E-H) Shoot true leaf formation recovered fully for all lovastatin concentrations applied [(E) 0 μM, (F) 0.1 μM, (G) 0.3 μM, (H) 1 μM] when transplanted to control medium at 4 DAG and grown for 10 d in control medium. (I) Whole ten-day-old seedlings. Note the formation of elongated lateral roots compared to mock. Scale bar: 1 mm (A-H), 5 mm (I).

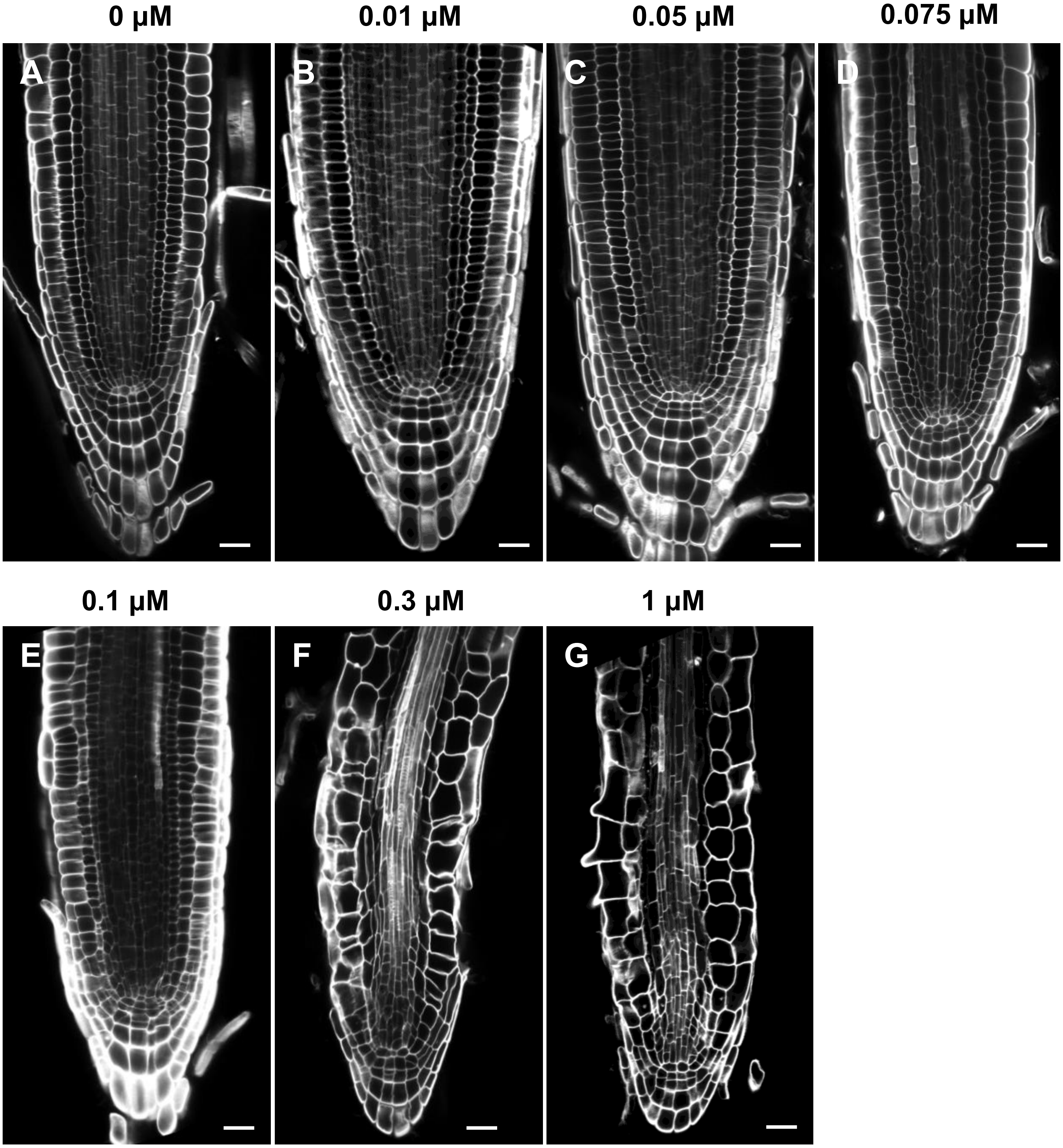


**Supplementary Fig. S4: Lovastatin affects root developmental zones but not the histological patterning, in a dose-dependent manner**. (A-G) Ten-day-old seedlings grown in various concentrations of lovastatin and counterstained with Calcofluor White. Note that under 0.1 µM the root apex does not show any obvious defect. At concentrations ≥ 0.1 µM meristem length is decreased dose-dependently and epidermis and cortex cells appear abnormally expanded. Scale bar 20 µm.

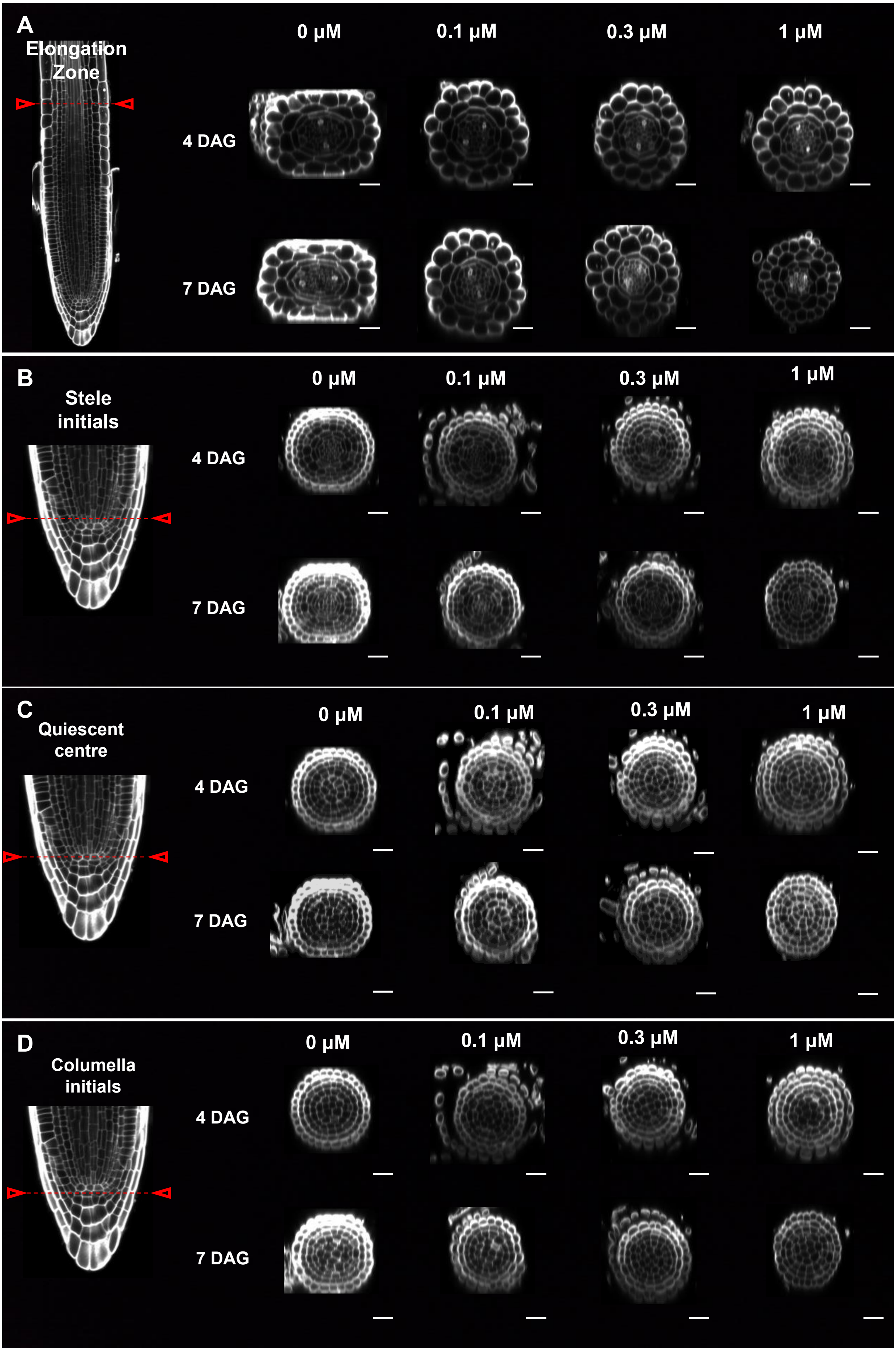


**Supplementary Fig. S5. Radial organization of primary roots is not affected by any lovastatin treatment applied.** Optical cross sections focusing on the (A) elongation zone, (B) stele initials, (C) QC and (D) columella initials of primary roots treated 4 DAG or 7 DAG with 0.1-1 µM lovastatin (arrows and dotted line mark the position of cross sections, longitudinal section demonstrates mock root tip). Scale bar: 20 µm.

**A**

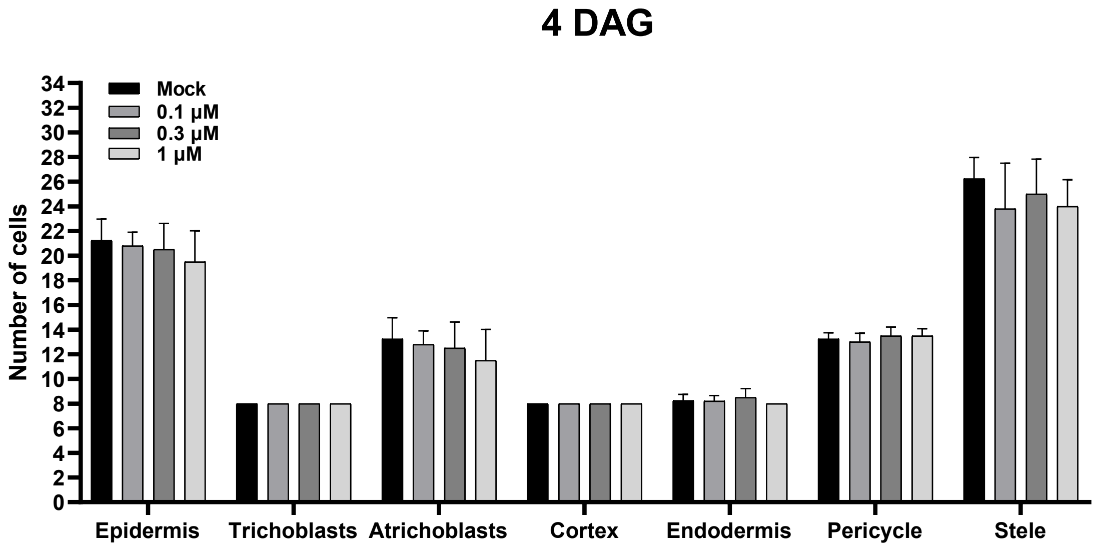


**B**

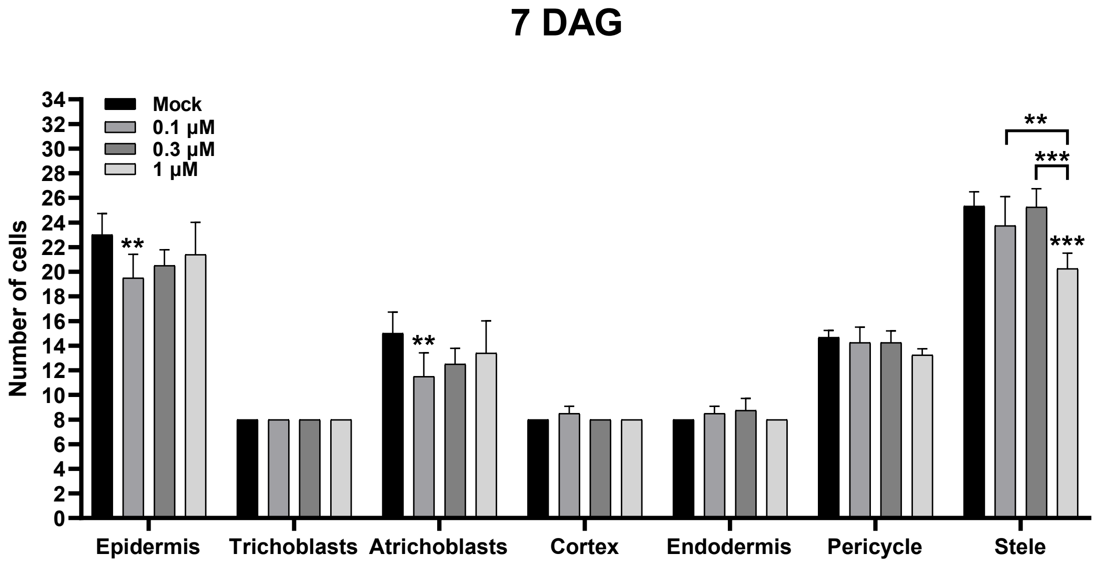


**Supplementary Fig. S6. Lovastatin did not affect total cell number across the root cell layers except atrichoblasts and stele at 7 DAG.** (A) Total cell number of lovastatin-treated seedlings 4 DAG did not show any variation compared to mock. (B) Lovastatin-treated seedlings 7 DAG demonstrated a slight decrease in atrichoblasts and stele cell number. For (A) and (B) cell number was calculated from the optical cross sections demonstrated in Supplementary Fig. S5. Two-way ANOVA combined with Tukey's for multiple comparison. Bar plots represent mean±standard deviation. N= 4 for all time points and concentration evaluated.

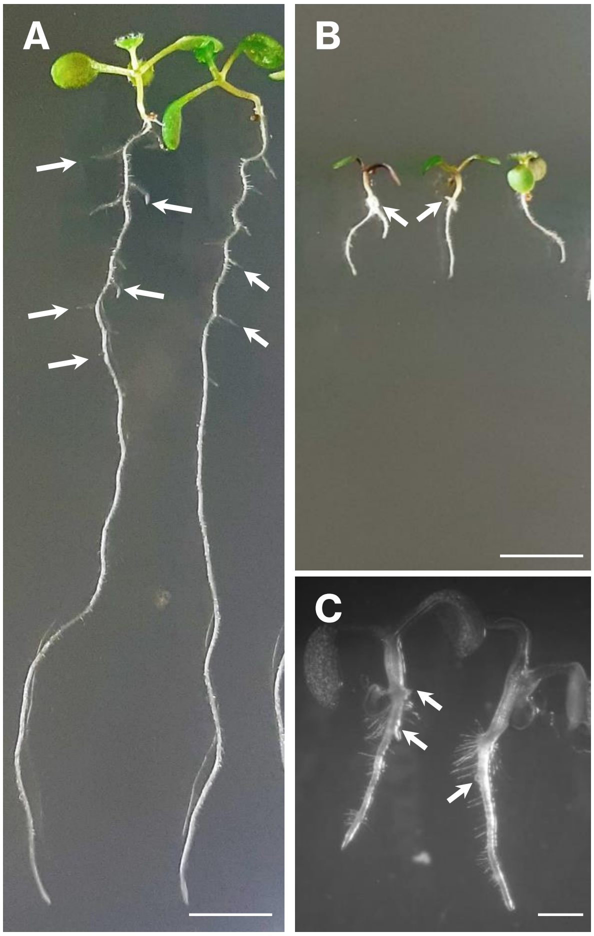


**Supplementary Figure S7. Inhibition of lateral root emergence after lovastatin treatment.** (A, B): Eight-day-old seedlings either untreated (A) or lovastatin-treated (B). Seedlings affected by lovastatin appear stunted with no true leaf development (C): Lovastatin-treated seedlings 14 DAG. Arrows indicate the lateral roots and adventitious roots (in treated seedlings). Note in (C) the abortive LR formation and the tumor-like shapes, also mentioned by Rodriguez-Conception *et al* (2004). Scale bar: 5 mm (A, B), 1 mm (C).

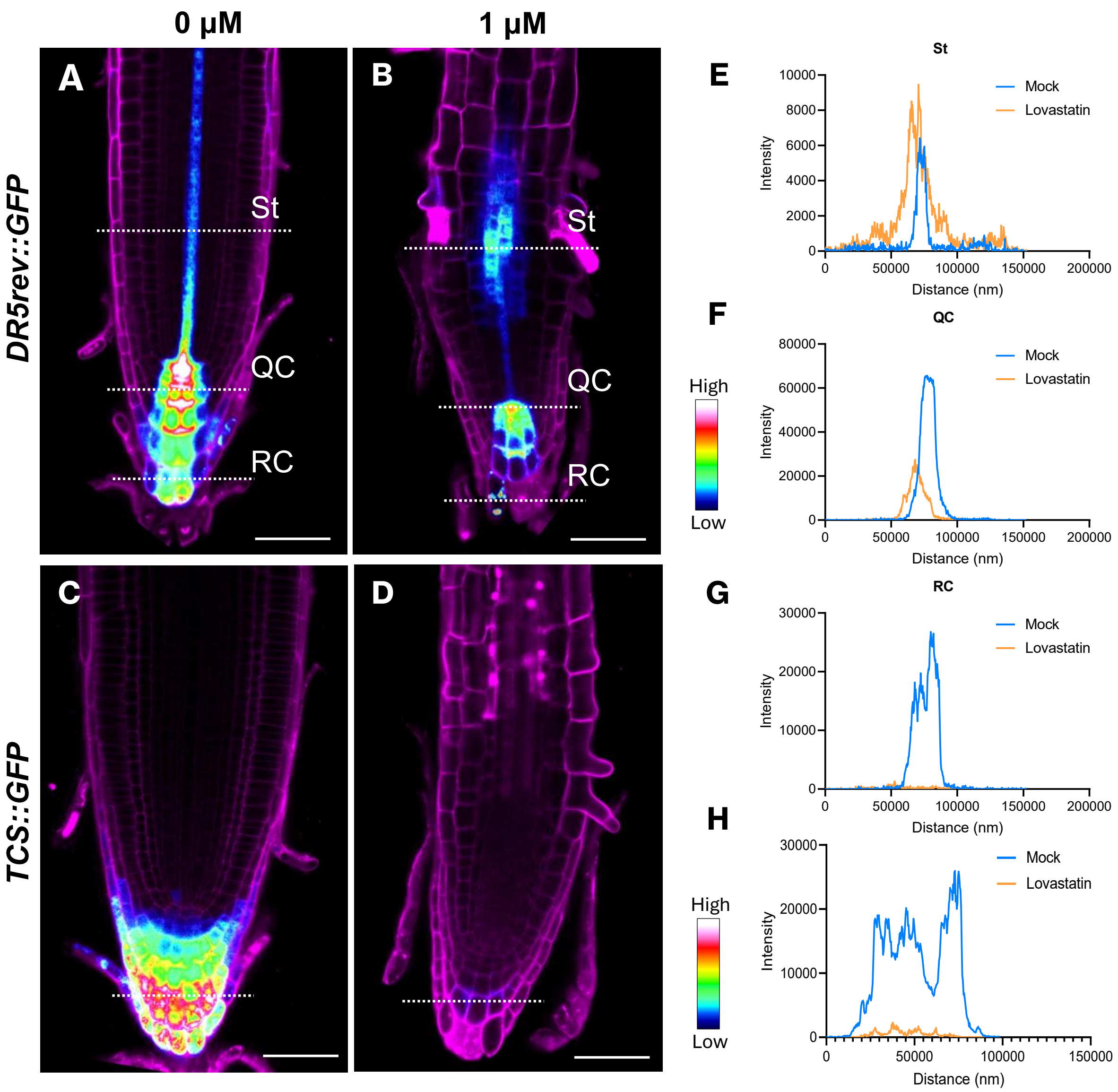


**Supplementary Fig. S8. Lovastatin treatment disturbed auxin gradient and almost completely abolished cytokinin at 7 DAG.** Propidium iodide stained *DR5rev::GFP* single CLSM sections of 7 d untreated (A) and 1 μM lovastatin-treated seedlings (B). Propidium iodide stained *TCS::GFP* CLSM images of 7 d untreated (C) and treated (D) roots. Intensity profiles at the line positions of *DR5rev::GFP* on root cap (RC) (E), QC (F) and stele (St) (G) were altered after lovastatin treatment. Intensity profile of *TCS::GFP* was almost abolished at the cytokinin collection region compared to untreated seedlings (H). At least 10 seedlings were evaluated for each condition. Scale bar: 50 μm.

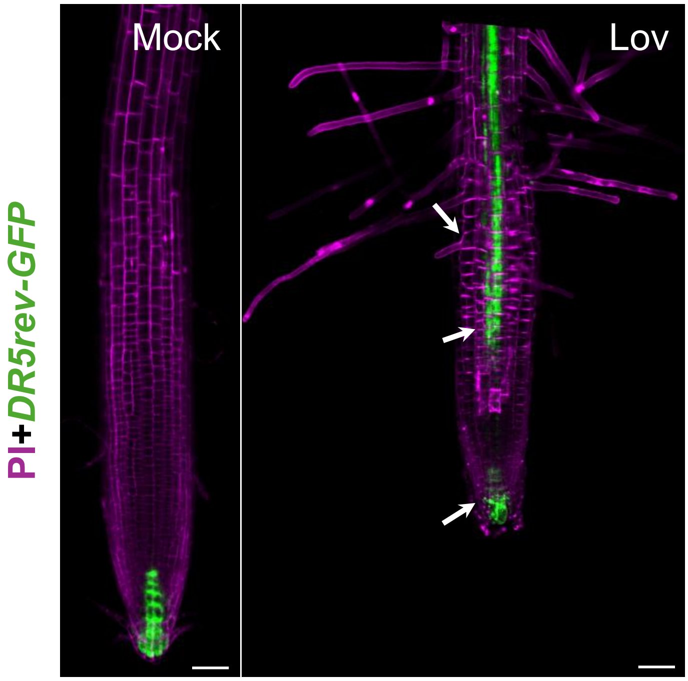


**Supplementary Fig. S9: Auxin gradient is significantly disrupted after lovastatin treatment.** Eight-day-old seedling germinated and grown in 1 µM lovastatin exhibits completely disrupted auxin maxima, in comparison to the mock, as revealed by *DR5rev-GFP*. Increased DR5-GFP signal along the root axis (arrows) indicating possibly a continuous signal for primordia initiation. Propidium iodide counterstaining. Scale bar 50 µm.

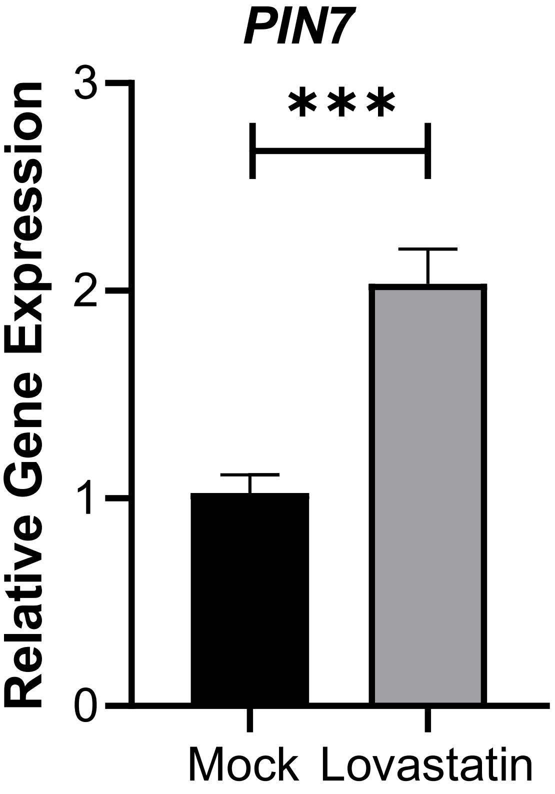


**Supplementary Fig S10. PIN7 increased gene expression after lovastatin treatment.** RT-qPCR analysis of PIN7 of mock (N=30) and 1 μM lovastatin-treated roots (N=90). Bar plots represent mean±standard error. Non-paramentic t-test applied. *** *p*<0.001.

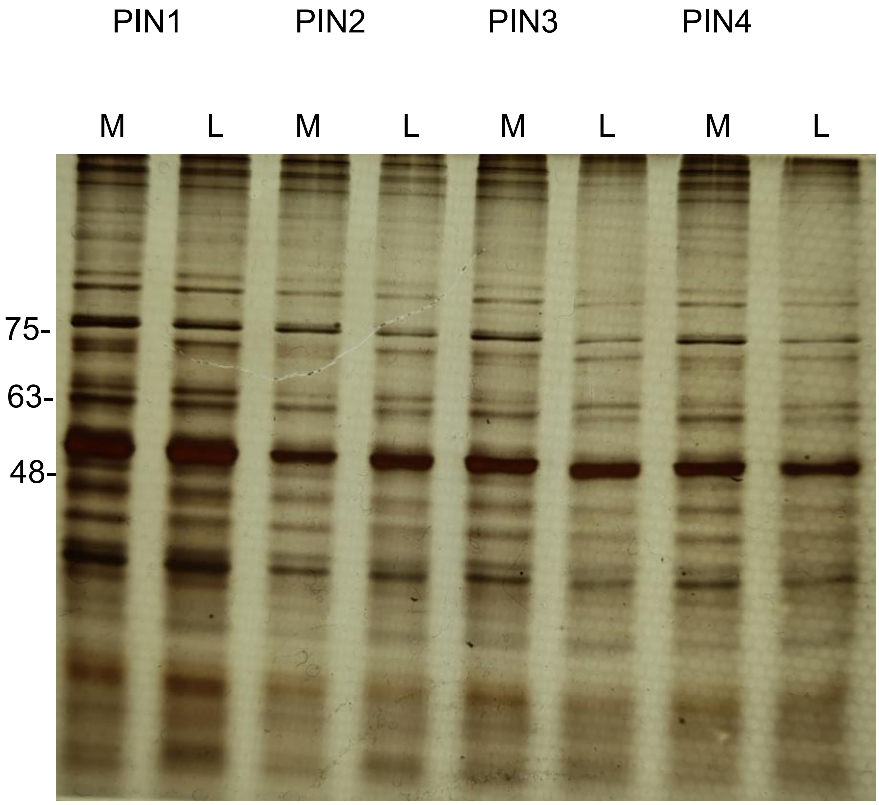


**Supplementary Figure S11:Total protein cell lysate profile of PINs 4DAG of roots untreated (M) and treated with 1 µM lovastatin (L).** Silver nitrate staining of untreated and treated roots of *PIN1::PIN1-GFP*, *PIN2::PIN2-GFP*, *PIN3::PIN3-GFP* and *PIN4::PIN4-GFP* reporter lines. 20 µg of total protein lysate were loaded in each lane.

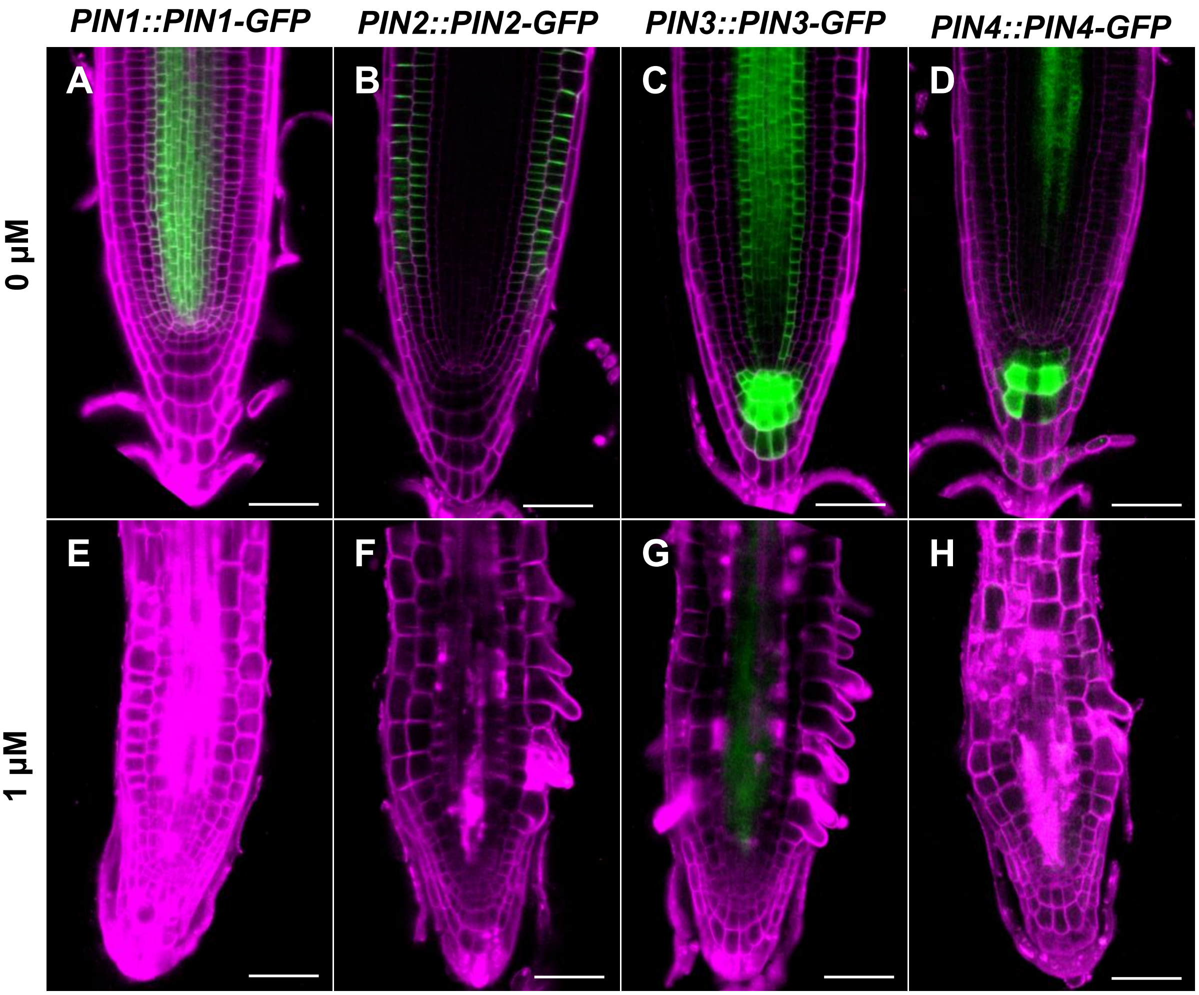


**Supplementary Figure S12. Plasma membrane localization of PINs is severely compromised at 7 DAG.** Single CLSM sections of propidium iodide stained 7 DAG of *PIN1::PIN1-GFP* (A, E), *PIN2::PIN2-GFP* (B, F), *PIN3::PIN3-GFP* (C, G) and *PIN4::PIN4-GFP* (D, H) reporter lines. At least 10 seedlings were used for each condition. Scale bar: 50 µm.

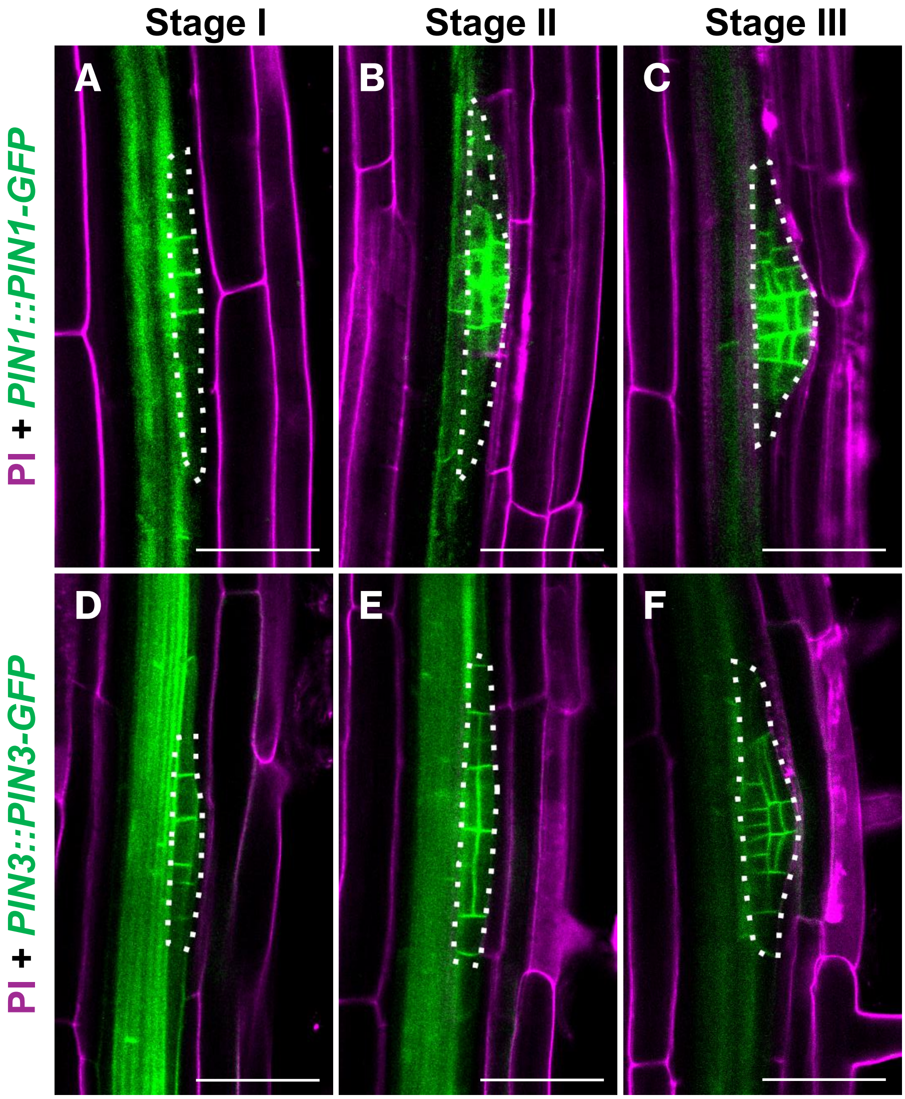


**Supplementary Fig. S13. Expression of PIN1-GFP and PIN3-GFP at early stages of lateral root development in mock seedlings.** (A) At the first stage PIN1-GFP is localized at the anticlinal faces of the initial cells. (B) At stage II PIN1-GFP signal is predominant at the periclinal faces of newly formed cells. (C) At stage III PIN1-GFP gradually becomes polarized towards the forming lateral root. (D-F) PIN3-GFP signal shows similar localization as PIN1-GFP but with more specific plasma membrane localization. Note that diffuse PIN1-GFP and PIN3-GFP signal can be detected in stele. Cell walls are counterstained with propidium iodide. Scale bar: 50 µm.
