## SupplementaryTables for "Effects of lovastatin on auxin transport and root development in *Arabidopsis thaliana*"

Supplementary Data

**Supplementary Table S1.** Reporter lines used in this study.

| Reporter line | NASC ID | Reference |
|---|---|---|
| *DR5rev::GFP* | N9361 | Benková et al. (2003) |
| *TCS::GFP* | - | Müller and Sheen (2008) |
| *PIN1::PIN1-GFP* | N9362 | Benková et al. (2003) |
| *PIN2::PIN2-GFP* | - | Xu and Scheres (2005) |
| *PIN3::PIN3-GFP* | - | Ioio et al. (2008) |
| *PIN4::PIN4-GFP* | N9576 | Vieten et al. (2005) |

**Supplementary Table S2.** Primers used in this study.

| Name | Sequence (5'-3') |
|---|---|
| KV139 PIN1 F | CGTTTGTGTTTGCCAAACAGT |
| KV140 PIN1 R | AACAAAGTCCCTGTGTTTTGGT |
| KV141 PIN2 F | TGCCAACGATAATGAGTGGA |
| KV142 PIN2 R | ATTTTCCGCACGCAATAATC |
| KV143 PIN3 F | GAGCACCTGACAACGATCAAG |
| KV144 PIN3 R | TCCACTTGCTGGATGAGCTAC |
| KV145 PIN4 F | TGCTAAGGAGATTCGGATGG |
| KV146 PIN4 R | AAGACCGCCGATATCATCAC |
| KV147 PIN7 F | TTCATCCCGCAATCTTGAGT |
| KV148 PIN7 R | ATCCTCTTCAGCCAAGCAGA |
| KB276 At4G26410 F | GAGCTGAAGTGGCTTCCATGA |
| KB277 At4G26410 R | GGTCCGACATACCCATGATCC |
